# Cell response to extracellular matrix energy dissipation outweighs rigidity sensing

**DOI:** 10.1101/2022.11.16.516826

**Authors:** Carla Huerta-López, Alejandro Clemente-Manteca, Diana Velázquez-Carreras, Francisco M. Espinosa, Juan G. Sanchez, Pablo Sáez, Álvaro Martínez-del-Pozo, María García-García, Sara Martín-Colomo, Andrea Rodríguez-Blanco, Ricardo Esteban-González, Francisco M. Martín-Zamora, Luis I. Gutierrez-Rus, Ricardo García, Pere Roca-Cusachs, Alberto Elosegui-Artola, Miguel A. del Pozo, Elías Herrero-Galán, Gustavo R. Plaza, Jorge Alegre-Cebollada

## Abstract

The mechanical properties of the extracellular matrix (ECM) determine cell differentiation, proliferation and migration through mechanoresponsive proteins including YAP. However, how different mechanical signals cooperate, synergize or compete to steer cell behavior remains poorly understood. Here, we have examined competition between the two major ECM mechanical cues, i.e. rigidity, which activates cell mechanosensing, and viscous energy dissipation, which reduces stiffness blunting cell mechanotransduction. To trigger competition, we have engineered protein hydrogels allowing concomitant modulation of stiffness and viscosity by mechanisms characteristic of native ECM. Culturing cells on these hydrogels, we have found that substrate energy dissipation attenuates YAP mechanosensing prevailing over stiffness cues. Hampered YAP activation on more dissipative substrates correlates with faster actin flow and smaller focal adhesions. Mechanistically, inhibition of actomyosin contractility reverses the outcome of the competition between rigidity and energy dissipation. Our results highlight the dominating contribution of substrate viscosity to the biology of the cell.

## Introduction

The extracellular matrix (ECM) is a protein hydrogel that provides support to cells in biological tissues. Structural and biochemical signals from the ECM are essential for cell homeostasis, adhesion, migration, proliferation, and differentiation ^1^. Notwithstanding the essential contribution of these classical cues to the biology of the cell, it is now well established that the mechanical properties of the ECM also impact extensively the activity and fate of cells ^2, 3^. A prime example is substrate stiffness ^2, 4^, which, when dysregulated, is a causative factor of aging ^5^ and diseases such as fibrosis and cancer ^6, 7^. Considering the pivotal role of ECM stiffness, it is tempting to hypothesize that other fundamental mechanical properties of the substrate, such as viscosity, can similarly steer cell behavior. Indeed, biological tissues and the ECM are viscoelastic materials that show substantial time-dependent drops in stiffness under strain, a property typically referred to as viscous energy dissipation ^8^. Remarkably, substrate energy dissipation has been observed to limit cell mechanosensing opposing the effects of rigidity ^9, 10^. However, how cells integrate and react to contradicting mechanical inputs remains largely unexplored. For instance, it is unknown whether energy dissipation can prevail over rigidity cues even when the apparent stiffness sensed by cells remains high ^11^.

The response of cells to the mechanical properties of the ECM has been studied using designer ECM-mimetic polymer hydrogels that are used as substrates for cell growth ^8, 12^. To limit confounding factors, optimal strategies achieve specific mechanical modulation while preserving non-mechanical features of the substrate that can also influence cell behavior, including pore size and mesh concentration ^13^. The use of ECM mimetics as cell substrates has been key to uncover mechanosensing and mechanotransduction systems deployed by cells to process mechanical inputs ^14^. In this regard, a central mechanism involves nuclear translocation of the transcriptional coactivator YAP, which constitutes a major proxy of the mechanobiological state of cells and is triggered when cells grow on stiff substrates ^15, 16^.

To investigate competition between substrate stiffness and energy dissipation in cell mechanosensing, we have concomitantly modulated both properties in protein hydrogels whose viscoelasticity stems from force-induced polypeptide conformational changes similarly to native ECM ^17^. Using these protein hydrogels as ECM mimetics, we have shown that actomyosin-based sensing of substrate energy dissipation leads to blunted YAP mechanosensing even when cells grow on stiff substrates.

## Results

### Tuning viscoelasticity of protein matrices

In a strained protein matrix such as the ECM, mechanical forces are directly transmitted to its protein building blocks ^18, 19^. When under force, random coil polypeptide regions easily adapt their extension to the external force in an elastic manner, while folded domains undergo reversible force-induced unfolding and extension transitions that provide energy dissipation ^20^ (**Fig. 1A**). Hence, proteins containing high proportion of random coil are soft and non-dissipative, whereas those rich in folded domains, particularly β structures, are stiffer and dissipate more energy. We reasoned that building on subtle sequence modifications that impact protein mechanics we could produce matrices with different viscoelastic properties but preserved structure and chemical composition. With that aim, we engineered polyprotein building blocks that contain eight repetitions of the 89-amino-acid-long immunoglobulin-like (Ig) domain I91 of titin ^21^. Key to our strategy, I91 domains have a single Tyr residue (Tyr9), which can be used to chemically crosslink soluble I91 polyprotein building blocks into hydrogel matrices using a [Ru(II)(bpy)_3_]^2+^-mediated photochemical reaction ^22, 23^.

**Fig. 1:**
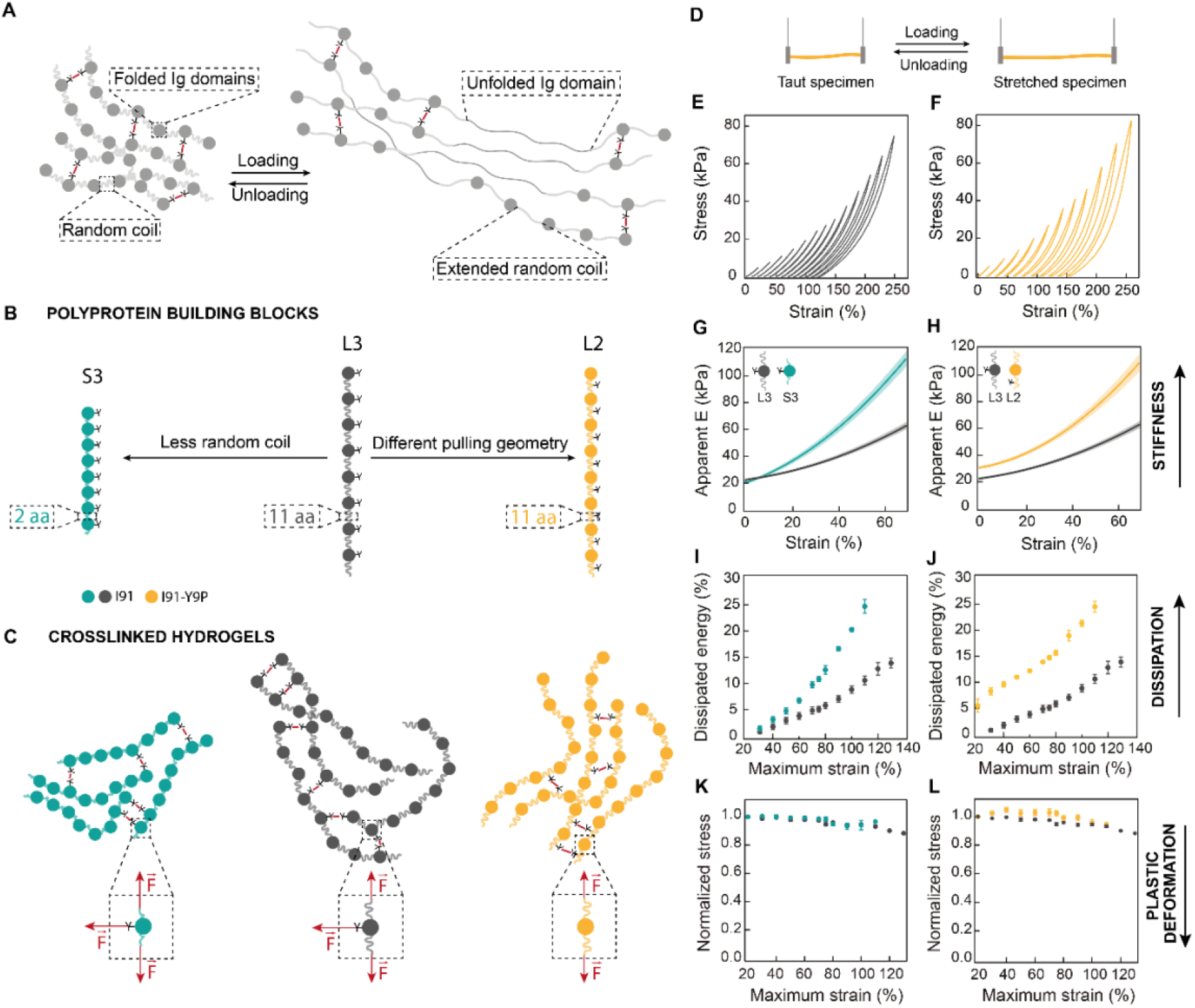
Engineered protein hydrogels with different viscoelastic properties. **(A)** Representation of a protein hydrogel matrix covalently crosslinked through tyrosine residues. Building blocks include Ig domains and random coil regions. Under uniaxial traction force, random coil regions extend elastically and Ig domains unfold, which dissipates energy. **(B)** Schematics of the polyprotein building blocks used in this study. Ig I91 domain was engineered into octameric repeats separated by 11 amino-acid-(aa)-long random coil linkers (L3, dark grey). Two more constructs were designed from L3 building blocks. First, S3 building blocks (teal) were produced by shortening the linkers between Ig domains to only 2 amino acids. To build the L2 building block (mustard), tyrosine residues were mutated out from Ig domains and included as one of the 11 amino acids in the random coil linkers. **(C)** Schematic representation of the network structure of I91 matrices. Crosslinking takes place at tyrosine positions, leading to different mechanical force distributions (dashed insets). **(D)** Loading and unloading stress-strain tests used to determine apparent elastic moduli, dissipated energy and plasticity of protein hydrogels. **(E, F)** Stress-strain analysis of L3 (E) and L2 (F) hydrogel matrices pulled at 5 mm/s. For clarity, stress–strain curves are offset in the x-axis. Actual stress–strain curves can be found in **Supplementary Fig. 5**. Curves were measured immediately one after another from low to high strains. **(G, H)** Strain dependency of apparent <u>elastic</u> moduli obtained in stress-strain experiments to 75 % maximum strain. Lines represent average values and shaded areas are standard error of the mean (SEM). **(I, J)** Dissipated energy is measured in stress-strain experiments to different maximum strains. **(K, L)** Normalized stress at 15% strain for consecutive stress-strain curves to different maximum strains. Data (mean ± SEM) are obtained from 4 specimens of S3 and L3 hydrogels and 6 specimens of L2 hydrogels produced in independent crosslinking reactions from 2 protein purification batches.

Since our goal was to trigger competition between substrate stiffness and energy dissipation, we aimed at producing protein hydrogels with concomitant changes in both mechanical properties. Our first strategy was to reduce the fraction of random coil structure by shortening the linkers between the I91 domains, which we hypothesized would lead to increased stiffness and energy dissipation. To this end, we produced two polyproteins in which the I91 domains are connected through polypeptide linkers of 2 and 11 amino acids (polyproteins S3 and L3, respectively; **Fig. 1B, Supplementary Fig. 1A**,**B, Supplementary Notes 1, 2**). The nomenclature refers to the length of the linkers between domains (S: short; L: long) and the number of pulling force vectors (2 or 3) applied to the domains, as explained below. We also explored if we could achieve equivalent modulation of viscoelasticity by exploiting the well-known influence of pulling geometry on protein nanomechanics ^24, 25^. For this second strategy, we produced polyprotein L2, in which the crosslinking tyrosine is mutated out from I91 and placed within the 11 amino-acid-long linker connecting I91 domains (**Fig. 1B, Supplementary Fig. 1A**,**C, Supplementary Note 3**). When strain is applied to L2 hydrogels, I91 domains experience only Nt-to-Ct pulling force (2 force vectors); while in S3 and L3 materials, two additional pulling geometries through crosslinked Tyr9 residues are possible for a total of 3 force vectors (**Fig. 1C**). Circular Dichroism (CD) and Differential Scanning Fluorimetry (DSF) showed that the overall β-rich structure and thermal stability of the I91 domain in the polyprotein building blocks are not affected by the length of the linkers, while mutation Y9P leads to lower thermal stability in agreement with previous observations ^26^ (**Supplementary Figs. 2, 3**).

We cast S3, L3 and L2 hydrogels to ∼13-mm-long and 0.506-mm-diameter cylinders using the same protein concentration and crosslinking conditions. We characterized the mechanical properties of the resulting hydrogels using uniaxial traction stress-strain tests (**Fig. 1D-L**), which show strain-stiffening and hysteresis between loading and unloading cycles that are typical of the ECM (**Fig. 1E,F**) ^8^. When comparing S3 and L3 specimens, which differ in the proportion of random coil regions, we found that the apparent elastic modulus (E) at zero strain of both hydrogels is ∼20 kPa. As strain increases, S3 hydrogels become pronouncedly stiffer than L3 counterparts, in agreement with the presence of shorter linkers in S3 building blocks (**Fig. 1G**). To extract information about viscous properties of the hydrogels, we determined how much energy is dissipated during loading and unloading stress-strain cycles. We found that S3 hydrogels are more dissipative than L3 counterparts, especially at medium and large strains, as expected from the higher contribution of unfolding transitions in S3 hydrogels (**Fig. 1I**). We obtained similar conclusions from indentation experiments using Atomic Force Microscopy (AFM) (**Supplementary Fig. 4**). Since plastic deformation can modify cell mechanosensing leading to convoluted results ^8, 27^, we verified that all loading stress-strain curves in these experiments overlap to a great extent (**Supplementary Fig. 5**). Indeed, the stress reached at 15 % strain for every curve is always >90% of the initial value (**Fig. 1K**), suggesting that plastic deformation in S3 and L3 hydrogels is marginal even at recovery times shorter than 5 seconds. Regarding our second strategy to tune viscoelasticity, we found that L2 hydrogels are also stiffer (**Fig. 1H**) and more dissipative than L3 counterparts (**Fig. 1J**). Similar to S3 and L3, L2 hydrogels show very little plastic deformation, if any (**Fig. 1L, Supplementary Fig. 5**).

We checked if mechanical modulation in S3, L3 and L2 hydrogels is achieved in the absence of non-mechanical alterations. Regarding chemical composition, the three hydrogels show overlapping infrared spectra in the amide I region (1700-1600 cm^-1^), which presents a maximum at 1628 cm^-1^ in agreement with the high β-structure content of the I91 domain (**Supplementary Figs. 1B,C, 6A**) ^28^. The thermal stability of I91 in the context of hydrogels, as determined by DSF, is lower than in soluble building blocks, although highly similar in the three materials (**Supplementary Fig. 3**). All hydrogels also show indistinguishable ultrastructure (**Supplementary Fig. 6B**). Finally, we examined crosslinking density using acid hydrolysis followed by amino acid analysis (**Supplementary Fig. 6C-F**). As it can be expected from identical crosslinking reaction conditions, tyrosines disappear to the same extent in the three hydrogels (**Supplementary Fig. 6D,E**). However, we detected less dityrosine products in L2 hydrolyzates (**Supplementary Fig. 6F**), raising the possibility that crosslinking in L2 matrices could be somehow hindered.

In summary, characterization of our engineered protein hydrogels indicates that S3 and L2 matrices are stiffer and more dissipative than L3 counterparts, and that these distinct mechanical properties occur alongside largely preserved structural and chemical properties. Hence, we set out to investigate how increased stiffness and energy dissipation in S3 and L2 hydrogels compete to drive cell behavior.

### Substrate energy dissipation blunts YAP activity

To examine competition between substrate stiffness and energy dissipation in cell mechanosensing, we cultured immortalized retinal pigment epithelial (RPE-1) cells on S3, L3 and L2 matrices and determined localization of YAP. Cells show normal attachment, morphology and global metabolic activity in all conditions (**Supplementary Fig. 7**). We also included control cultures on elastic, non-energy-dissipating polyacrylamide (PAAm) hydrogels, in which we observe the expected nuclear localization of YAP on stiff (∼20 kPa), but not on soft (∼2.8 kPa), substrates ^15^ (**Fig. 2A,B**). YAP nuclear localization on stiff PAAm correlates with increased expression of YAP target genes ANKRD1 and CTGF ^29^ (**Supplementary Fig. 8**) and cell spreading (**Fig. 2C**). Regarding cells cultured on protein matrices, we observe the most intense nuclear YAP staining in cells cultured on L3, the softest and least dissipative (**Fig. 2A,B**). Increased YAP nuclear localization is accompanied by higher expression levels of YAP target genes, especially when compared to S3 matrices (**Supplementary Fig. 8**), and increased cell spreading (**Fig. 2C**). Higher YAP nuclear localization in L3 matrices correlated with formation of larger focal adhesions, a situation also observed in elastic hydrogels ^30^ (**Fig. 2D,E**). Using AFM indentation measurements, we confirmed that S3 matrices remain stiffer and more dissipative than L3 counterparts after cell culture (**Supplementary Fig. 9**). Supporting the generality of our observations, we obtained similar patterns of YAP activation and cell spreading in mesenchymal stem cells (**Supplementary Fig. 10)**. Our data showing lower YAP nuclear translocation on S3 and L2 matrices indicate that ECM energy dissipation blunts YAP activation. Since energy dissipation leads to reduced effective stiffness, rigidity sensing mechanisms may suffice to explain our results if the actual mechanical stress sensed by cells was the highest in L3 matrices. To test this possibility, we stretched S3 and L3 hydrogels to strains between 10-50%, which is the range typically induced by cells ^31^, and monitored the resulting stress for 3 hours. Results show that S3 hydrogels generate higher stress than L3 counterparts at all times despite being more dissipative (**Fig. 3A-C**). Indeed, although the ratio between the stress generated by S3 and L3 hydrogels decreases with straining time as expected, S3 hydrogels are predicted to remain stiffer even after 24 hours under strain (**Fig. 3D**). Altogether, our results show that energy dissipation leads to reduced nuclear translocation of YAP outweighing stiffness cues.

**Fig. 2:**
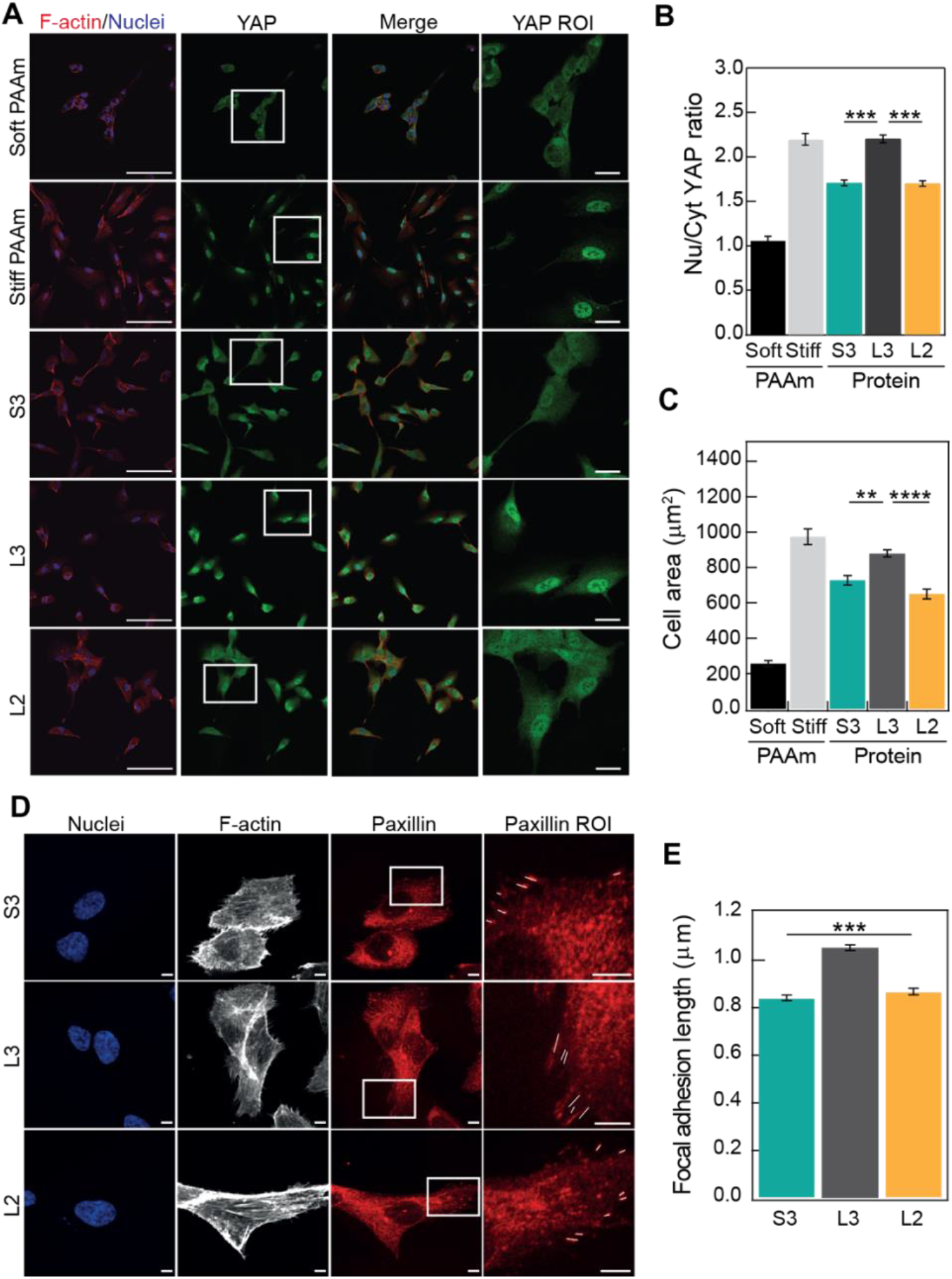
Blunted YAP activity in cells grown on more dissipative substrates. **(A)** Confocal immunofluorescence images of YAP localization in RPE-1 cells grown on elastic stiff or soft PAAm substrates and on different I91 matrices. F-actin was stained with alexa647-conjugated phalloidin (red; first column), and nuclei were stained with DAPI (blue in merged images; first and third column). YAP was labelled with alexa488 conjugated antibody in green (second column). The fourth column shows zoomed views of the YAP ROI (boxed in white in the YAP column). Scale bars are 100 μm (first column) and 20 μm (fourth column). **(B)** Quantification of nuclear versus cytoplasmic YAP distribution. **(C)** Cell spreading quantified as cell area. A minimum of n=30 cells per condition were quantified in a total of 4 independent experiments. **(D)** Confocal immunofluorescence images of focal adhesions in RPE-1 cells. Nuclei were stained with DAPI (first columns) and F-actin was stained with alexa647-conjugated phalloidin (second column). Paxillin in focal adhesions was labelled with alexa568-conjugated antibody in red (third column). The fourth column shows zoomed views of the paxillin ROI (boxed in white in the paxillin column). Scale bars are 5 μm. **(E)** Focal adhesion length quantification. A minimum of n=30 cells per condition were quantified in a total of 3 independent experiments. Data are presented as mean ± SEM.

**Fig. 3:**
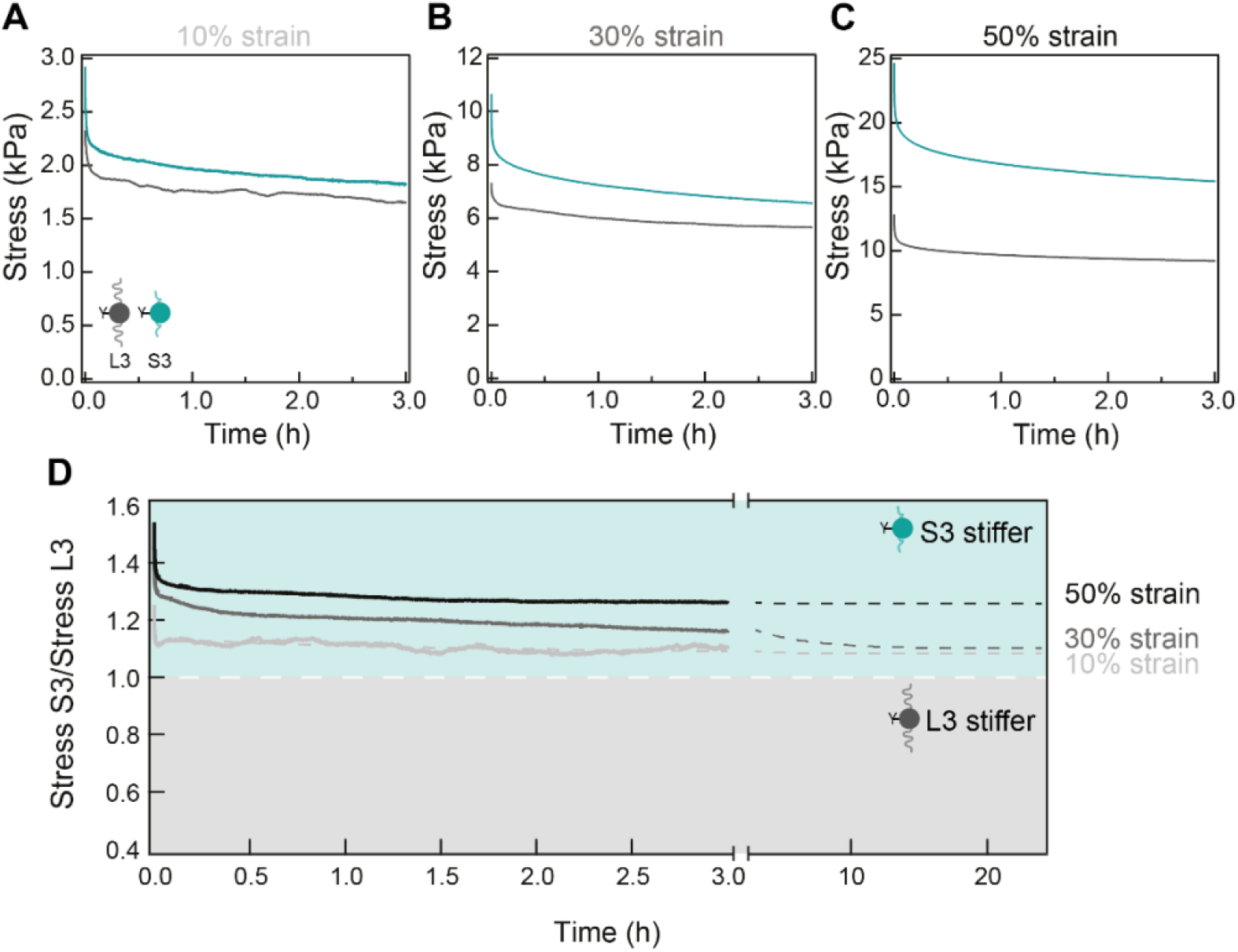
S3 hydrogels remain stiffer than L3 counterparts under long-duration strains. Stress-relaxation of S3 (teal) and L3 (dark grey) hydrogels for 3 hours at a **(A)** 10 % **(B)** 30% and **(C)** 50 % strain. **(D)** Ratio of the stress generated by S3 and L3 hydrogels in the relaxation experiments. Ratios above 1 indicate S3 hydrogels are stiffer than L3 counterparts. Grey dashed lines are double-exponential fits to the experimental data. Results were obtained with three L3 and three S3 specimens coming from three crosslinking reactions and two different protein purifications.

### Energy dissipation sensing requires actomyosin contractility

Cells probe the mechanical properties of the ECM through dedicated contractile and adhesive molecular machinery. The molecular clutch model, which is based on the integrated force response of the ECM, integrins, adaptor proteins and the actomyosin cytoskeleton, can explain cell mechanotransduction on elastic substrates ^30^ (**Fig. 4A**). According to this model, the force generated by myosin contractility and actin polymerization propels actin retrograde flow from the cell membrane towards the nucleus, pulling on integrin-ECM molecular clutches. Above a stiffness threshold, the clutches experience high forces that result in the mechanical unfolding of the adaptor protein talin, which induces the recruitment of additional adaptor proteins. This chain of events reinforces adhesion, reducing actin retrograde flow and leading to YAP translocating to the nucleus ^30^. Adaptation of the molecular clutch to purely viscous substrates predicts that energy dissipation opposes clutch engagement, resulting in increased actin retrograde flow and reduced YAP nuclear translocation ^32^. In agreement with this model, we detect faster actin retrograde flow in the more dissipative S3 and L2 matrices (**Fig. 4B,C**).

**Fig. 4:**
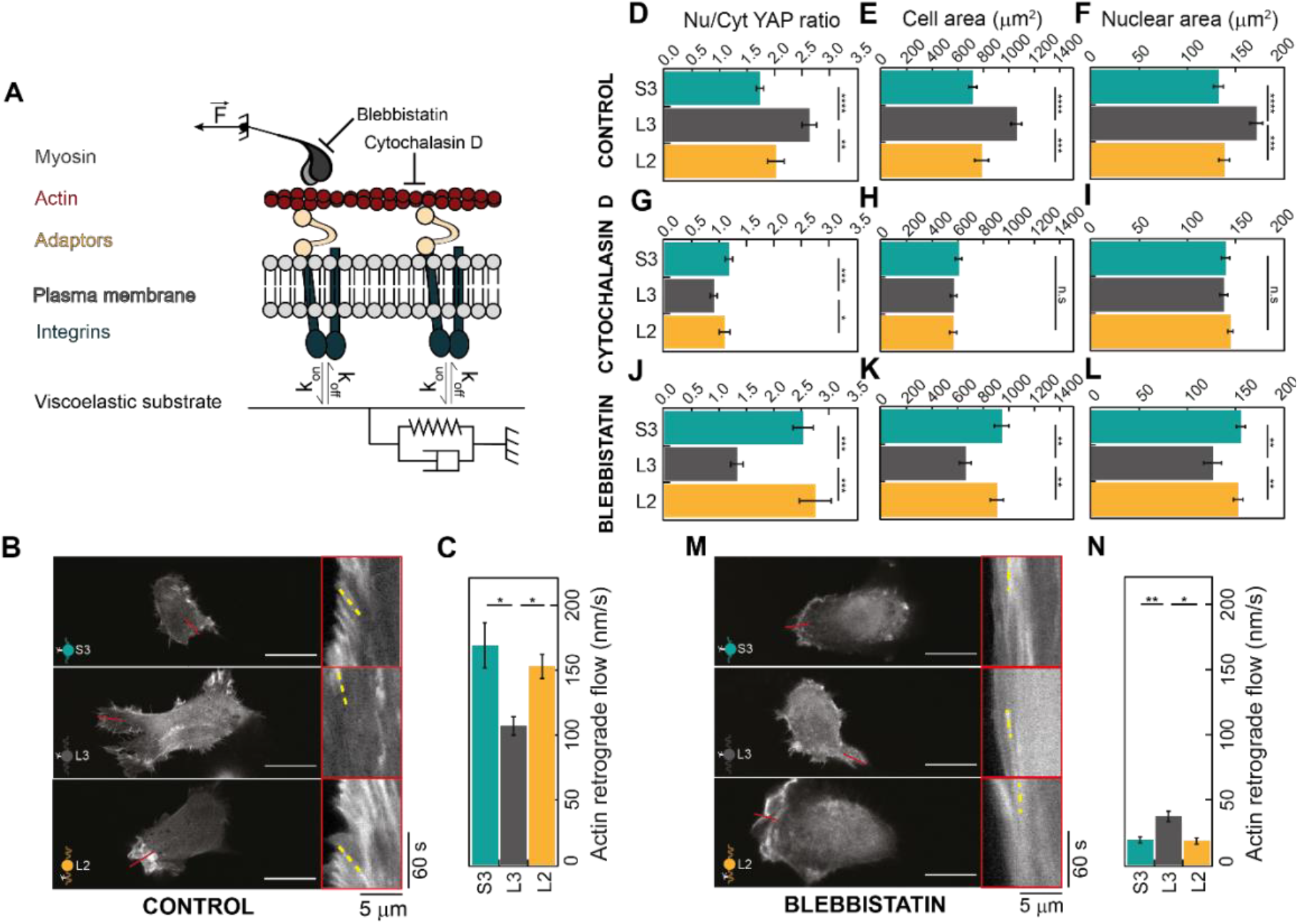
Energy dissipation sensing requires actomyosin contractility. **(A)** Mechanical communication between a viscoelastic ECM and cell cytoskeleton takes place through molecular clutches of integrins and adaptor proteins. Blebbistatin and cytochalasin D treatment inhibits myosin activity and actin polymerization, respectively. **(B)** Actin retrograde flow of cells seeded on S3, L3 and L2 matrices was measured in LifeAct-GFP time-lapse acquisitions. Representative static images are shown on the left (20 μm scale bars). Kymographs on the right show actin movement towards the cell center measured in regions of interest indicated by red lines. Representative actin trajectories are highlighted in yellow. **(C)** Quantification of actin retrograde flow from at least 30 measurements per condition. **(D-F)** Quantification of nuclear versus cytoplasmic YAP distribution, cell spreading and nuclear area of cells seeded on S3, L3 and L2 matrices. **(G-I)** Quantification of YAP translocation to the nucleus, cell spreading and nuclear area of cells seeded on S3, L3 and L2 matrices and treated with 1 μM cytochalasin D. **(J-L)** Quantification of YAP translocation to the nucleus, cell spreading and nuclear area of cells seeded on S3, L3 and L2 matrices and treated with 10 μM blebbistatin. A minimum of 40 cells coming from 3 independent experiments were analyzed for quantifications in D-L. **(M)** Actin retrograde flow of cells seeded on S3, L3 and L2 matrices and treated with 10 μM blebbistatin. Representative static images are shown on the left (20 μm scale bars). Kymographs on the right show actin movement towards the cell center measured in regions of interest indicated by red lines. Representative actin trajectories are highlighted in yellow. **(N)** Quantification of actin retrograde flow from at least 30 measurements per condition.

We investigated how individual components of the clutch contribute to the response of cells to competing stiffness and energy dissipation of the ECM. In agreement with results in **Fig. 2B,C**, higher YAP activity on RPE-1 cells cultured on L3 hydrogels correlates with increased cell and nuclear areas in control experiments (**Fig. 4D-F, Supplementary Fig. 11**). In cells treated with cytochalasin D, an actin polymerization inhibitor, nuclear YAP levels decrease regardless of the substrate used (**Fig. 4D,G, Supplementary Fig. 11**). Under these conditions, differences in YAP localization do not correlate with cell and nuclear areas although they appear to follow the stiffness of the substrate (**Fig. 4G-I**). This effect is more marked when cells are incubated with myosin inhibitor blebbistatin ^30^, which leads to YAP activities that correlate with cell and nuclear areas (**Fig. 4J-L, Supplementary Fig. 11**). Remarkably, treatment with blebbistatin results in higher YAP nuclear localization in S3 and L2 matrices, but lower in L3, as compared to control conditions (**Fig. 4D,J**). Actin retrograde flow in the presence of blebbistatin correlates with YAP activation state as in control cells, although myosin inhibition leads to lower absolute flow rates as expected (**Fig. 4B-D,J,M,N**) ^33^.

## Discussion

Mechanical signals are inextricably bound to tissue and cell homeostasis. Here, we have harnessed polypeptide mechanics to engineer protein ECM mimetics with tailored viscoelastic properties but preserved ultrastructure and chemical composition. By culturing cells on these matrices, we have uncovered that substrate energy dissipation in 2D cell cultures results in lower YAP nuclear translocation outweighing rigidity cues. Interestingly, our data show that inhibiting mechanisms of cell force generation leads to mechanotransduction that follows the rigidity of the substrate, pointing to their essential role in the integration of substrate stiffness and energy dissipation cues.

We propose that the molecular clutch conceptual framework is compatible with the observation that YAP nuclear translocation is lower in more dissipative substrates (**Fig. 2**), as well as with the inverted behavior we have found under actomyosin inhibition (**Fig. 4**). Since the stiffness of S3, L3 and L2 matrices is high, clutches could reach the force threshold for talin unfolding enabling reinforcement of cell mechanotransduction under the three control conditions ^30^. However, after reaching that point substrate energy dissipation would result in progressively lower effective force on talin. Since talin rapidly fluctuates between folded and unfolded states ^34^, this lower effective force could favor the folded state, leading to less reinforcement, smaller focal adhesions and reduced YAP nuclear translocation on more dissipative substrates. Under the lower myosin pulling rates induced by blebbistatin ^35^, the substrate stiffness requirement for talin unfolding increases ^30^ and is probably not reached by the softer L3 matrix, in agreement with the reduced YAP nuclear translocation we have detected when myosin is inhibited in cells growing on this substrate. For the stiffer S3 and L2 substrates, the force regime for talin unfolding could still be reached, although with a delay with respect to control experiments due to the lower myosin pulling rates. This delay would imply that S3 and L2 substrates dissipate more energy in the process of reaching the regime for talin unfolding than in control situation, and therefore have less energy left to dissipate by then. Due to this somewhat counterintuitive effect, the force experienced by talin in the unfolding regime would be higher when myosin is inhibited in S3 and L2 matrices, resulting in a more populated talin unfolded state, increased adhesion strength and more translocation of YAP to the nucleus.

Our results highlight how stiffness, albeit essential, may not be the main mechanical parameter of the ECM driving cell behavior ^9, 27, 36, 37^. Indeed, we have shown that the dissipative properties of the ECM can lead to cell responses that are unpredictable considering only substrate stiffness. Hence, we propose that characterization of the viscous properties of tissues, cells and the ECM is essential to fully apprehend the complex crosstalk between biology and mechanics. For instance, the reported negative correlation between ECM viscosity and tumor malignancy could be explained by the energy-dissipation-driven inhibition of YAP that we have detected, since YAP activation typically results in more aggressive tumors ^38^. Based on our results, it is conceivable that disruption of the homeostatic balance between substrate stiffness and energy dissipation can impact cell function negatively ^39^. It follows that cell response to viscoelasticity should be considered and integrated in holistic tissue engineering and regenerative medicine applications.

## Supporting information

HuertaLopez_SupplementaryInfo

## Acknowledgements

We thank CNIC’s Microscopy Facility for assistance with light microscopy experiments and the Spectroscopy and Nuclear Magnetic Resonance Core Unit at CNIO for access to CD instrumentation. We thank Sebastián Prieto (Sereetron), Daniel Jiménez and David Bartolomé (CNIC) for their support to configure the tensile tester, and Konrad Güth for providing hydrogel gripping tweezers. We thank Miguel Vicente Manzanares, Andrés Hidalgo, David de Sancho and all members of the *Molecular Mechanics of the Cardiovascular System* team for their feedback.

## Author contributions

CHL and JAC conceived the project. CHL, AMP, PS, MGG, RG, AEA, PRC, MAP, EHG, GP and JAC designed research. CHL, ACM, DVC, FME, JGS, AMP, ARB, REG, FMZ, LGR, SMC and GP executed experiments. CHL and ACM analyzed experiments. RG, MAP, PS, AEA, PRC, GP, JAC ensured funding. CHL and JAC drafted the manuscript with input from all authors.

## Data and materials availability

Supplementary information is available in the online version of the article. Materials and raw data are available from the corresponding author upon reasonable request.

## Materials and methods

### Polyprotein Engineering

The complementary DNA (cDNA) coding for titin I91-Y9P was a gift from Mariano Carrión-Vázquez (Instituto Cajal). Linker amino acids and the extra tyrosine needed for crosslinking were added by PCR (Polymerase Chain Reaction) using the primers in **Supplementary Table 1**. The cDNA coding for I91 was provided by Raúl Pérez-Jiménez (CIC nanoGUNE). cDNAs coding for octamers were produced following an iterative strategy of cloning using BamHI, BglII, and KpnI restriction enzymes, except for the I91_8_ cDNA to produce the L3 building block, which was synthesized by GeneArt (Thermo Fisher). cDNAs were inserted in the pQE80 expression plasmid (Qiagen, Valencia, California, USA) using BamHI and BglII restriction enzymes, and the resulting plasmids were verified by Sanger sequencing. Full polyprotein sequences can be found in **Supplementary Notes 1-3**. Polyproteins were expressed in *Escherichia coli* BLR (DE3). Briefly, fresh cultures (OD_600_ = 0.6–1.0) were induced with 0.4 mM (S3, L2) or 1 mM (L3) isopropyl β-D-1-thiogalactopyranoside for 3 h at 37 °C and at 250 rpm. Cells were lysed by a combination of tip sonication and passes through a French Press. Polyproteins were purified from the soluble fraction through Ni-NTA agarose affinity chromatography (HisTrap FF, 5 mL), carried out in a Fast Protein Liquid Chromatography system (GE Healthcare). Proteins were eluted in 50 mM sodium phosphate pH 7, 300 mM NaCl, 250 mM imidazole, and dialyzed against 50 mM sodium phosphate pH 7.5, 150 mM NaCl. Purified polyproteins were concentrated by filtration using 2 mL Amicon Ultra 4 tubes. Purity of samples was evaluated using sodium dodecyl sulfate-polyacrylamide electrophoresis gels (**Supplementary Table 1**). We obtained 30-40 mg polyprotein per liter of bacterial culture. Protein concentration was estimated from A_280_ measurements considering theoretical extinction coefficients (**Supplementary Table 2**). Protein samples were stored at 4 ºC with 0.1 % sodium azide.

### Circular dichroism

CD spectra were collected using a Jasco J-810 spectropolarimeter. All soluble polyproteins were tested in 50 mM sodium phosphate pH 7.5, 150 mM NaCl buffer in 0.1 cm-pathlength quartz cuvettes at ∼0.3 mg/mL protein concentration. Spectra were recorded at a scanning speed of 50 nm/min and a data pitch of 0.2 nm. Averaged final spectra are the result of four accumulations. The contribution of the buffer was subtracted from each protein spectrum, which was normalized by peptide bond concentration. To study thermal denaturation, the CD signal at 215 nm was monitored as temperature increased from 25 to 85°C at a rate of 30°C/h (0.5°C data pitch). Temperature control was achieved through a Peltier thermoelectric system.

### Protein hydrogel preparation

Hydrogel gelation was achieved following a [Ru(II)(bpy)_3_]^2+^ mediated photochemical crosslinking strategy targeting tyrosine residues ^18, 22, 23^. To this aim, 1 mM purified protein was mixed with 25% ammonium persulfate (APS) (Sigma-Aldrich) and 6.67 mM [Ru(II)(bpy)_3_]^2+^ (Sigma-Aldrich). To produce cylindrical hydrogels for uniaxial traction testing, the mix was loaded into polytetrafluoroethylene (PTFE) tubing (Cole-Parmer) (d_in_ = 0.022 inch, d_out_ = 0.042 inch) through the application of negative pressure using a 25 G needle attached to a syringe. A 2400 lm LED white light source, with an emission maximum of 452 nm, was placed directly below the tubing to irradiate the sample for 3 hours in a cabinet at 4°C (due to local heating, the resulting reaction temperature was measured to be

∼25°C). The crosslinked specimen was then taken out the PTFE tube and stored in 50 mM sodium phosphate pH 7.5, 150 mM NaCl buffer at 4°C. Hydrogels prepared following this procedure can be stored for several weeks without affecting their mechanical response. To produce disk-shaped hydrogels, polyproteins were crosslinked on round glass coverslips (d=10 mm) that had been plasma activated using a Plasma Cleaner (Harrick Plasma) for 10 minutes at high intensity. To this aim, 7.1 μL of the reaction mix were placed on a Gel Slick™ (Lonza)-coated glass slide, and then a plasma-activated coverslip was gently pushed against the protein solution. Slide-coverslip sandwiches were then transferred to the surface of the LED lamp and irradiated for 3 hours. After crosslinking, hydrogels were thoroughly washed with Phosphate-buffered saline (PBS), coated with 5 μg/mL of fibronectin (Sigma-Aldrich, F1141) for 1 hour at 37 ºC, then washed 3 times and preserved in PBS at 4 ºC no longer than 24 hours.

### Differential Scanning Fluorimetry

Melting curves were determined by mixing 10 μg of protein, either soluble or crosslinked hydrogels, with SYPRO Orange (ThermoFisher, S6650) following the manufacturer specifications. Samples were taken to a final volume of 25 μL with 50 mM sodium phosphate pH 7.5, 150 mM NaCl and were allowed to incubate with the dye for 1 hour at 4 ºC before being loaded into a 384-well qPCR plate (ThermoFisher, BC2384). The plate was then placed into a CFX384 Touch Real-Time PCR Detection System (BioRad) and melting curves were obtained raising the temperature from 10 to 95 ºC at 3 degrees per minute. SYPRO Orange fluorescence at 570 nm increases when proteins denature until proteins aggregate, which leads to a decrease in fluorescence signal. The global minimum of the first derivative of the melting curve indicates melting temperatures.

### Tensile Testing

Tests were done using a home-built tensile tester inspired by Saqlain and colleagues ^31^. The tester consists of a SI-H KG4A force sensor (World Precision Instruments) hooked to a BAM21-LC KG Optical Force Transducer Amplifier (World Precision Instruments) and a SMAC LCA25-025-35-6F linear actuator (SMAC Moving Coil Actuators) connected to a computer through a NIDAQ USB-6002 data acquisition system (National Instruments). The force sensor and linear actuator are both connected to a pair of stainless-steel SI-TM7 general-purpose mount tweezers used to grip the specimen. For tensile testing, cylinder shaped hydrogels were fixed at both ends by the tweezers. Tensile tests were done in 50 mM sodium phosphate pH 7.5, 150 mM NaCl buffer at room temperature. In stress-strain tests, strain was applied and then removed at a 5 mm/s rate, whereas in stress-relaxation measurements, the strain was held constant while the load was recorded as a function of time. Apparent E were calculated from stress-strain curves performed to 75% strain as the derivative of stress (tangent modulus). Dissipated energy was evaluated from stress-strain curves carried out at increasing strain values as the ratio of the hysteresis area between the loading and unloading stress-strain curves and the area below the loading stress-strain curve.

### Nanomechanical spectroscopy by Atomic Force Microscopy

AFM measurements were performed with a commercial instrument, JPK NanoWizard 3 (JPK Instruments AG, Berlin, Germany), mounted on an Axio Observer D1 inverted microscope (Carl Zeiss, Oberkochen, Germany). We used BL-AC40TS cantilevers. The force constant was calibrated using the thermal noise method (*k* =0.089 N/m). The optical sensitivity was 7.2 nm/V. The nominal values for the tip’s geometry, half-angle, height and radius, were, respectively, 18º, 7 µm and 8 nm.

Force-distance curves (FDC) were performed on 5 different positions of the hydrogel samples; 10 force-distance curves were acquired at each position. The FDCs were acquired by applying a triangular waveform at *v* = 10 µm/s with a data acquisition rate of 5 kHz. The tip’s indentation was stopped when the force reached 3 nN. The z-piezo displacement was 5 µm (closed-loop feedback). Mechanical parameters were obtained by fitting the FDCs curves with a 3D power law rheology model that has two parameters, a modulus *E*_0_ that relates to hydrogel stiffness, and an exponent γ that is linked to energy dissipation ^40^. Specifically, we have used the expressions for a conical tip assuming a semi-infinite sample thickness and fitting values were filtered according to the correlation coefficient *R* (≥0.95):

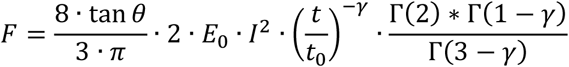

*F* is the force applied to the hydrogel, *I* is the indentation, *θ* is the cone half angle, and Γ the gamma function. The 3D power law rheology model determines the indentation differently for the approach (app) and retraction (ret) sections of the FDC,

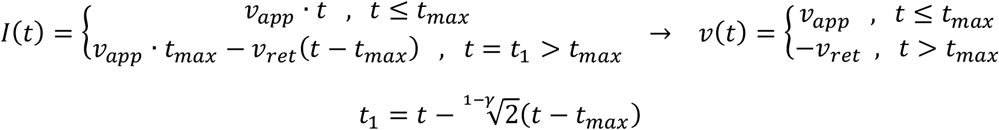

where *t*_max_ is the time at which maximum indentation is reached.

### Infrared spectroscopy

Attenuated total reflection (ATR) Fourier transform infrared (FTIR) spectra were obtained using a Nicolet iS5 spectrometer equipped with an iD5 ATR complement following published protocols ^41^. ATR-FTIR spectra were obtained from hydrogels pressed against the diamond window at constant pressure in 50 mM sodium phosphate pH 7.5, 150 mM NaCl. A background spectrum obtained while applying constant pressure to buffer was subtracted from the spectrum of each sample. All spectra were the result of averaging 32 measurements in the 550-4000 cm^-1^ range, with a 4 cm ^-1^ resolution.

### Scanning Electron Microscopy

I91 hydrogels in 50 mM sodium phosphate pH 7.5, 150 mM NaCl were snap frozen in liquid nitrogen. Frozen samples were quickly transferred to a freeze dryer (VirTis) and lyophilized for 24 hours. The resulting specimens were then fractured in liquid nitrogen. The surface of the samples was coated with 6.9 nm carbon using a Leica EM ACE600 coater before observation. Imaging of the fractured end of hydrogels was performed using a Zeiss Auriga Field-Emission Scanning Electron Microscope. To quantify pore size, images were processed in FIJI. An internal gradient with a radius of 15 pixels was applied to each image to better define the areas corresponding to pores. Afterwards, a local threshold was applied and the area of the thresholded pores was obtained using FIJI’s Analyze Particles plugin.

### Quantification of dityrosine crosslinks

For analysis of the amino acid content of soluble polyproteins and crosslinked hydrogels, samples containing 10 nmol of tyrosine were hydrolyzed in 0.2 mL 5.7 N HCl containing 0.1% phenol and 119 μM N-Leu as an internal standard in evacuated sealed tubes at 110°C for 24 hours. Samples were then dried and washed 3 times with milli-Q water. Subsequent amino acid analyses were performed on a Biochrom 20 automatic analyzer (Pharmacia) following the procedures recommended by the manufacturer.

### PAAm hydrogel preparation

PAAm matrices were prepared on glass coverslips as previously described ^42^. Briefly, the surface of the coverslips was treated with (3-Aminopropyl)triethoxysilane (APTES) (Sigma-Aldrich, 281778) to facilitate PAAm adhesion. To ensure the formation of a homogeneous PAAm layer on the glass surface, the mixture was sandwiched between the APTES-treated coverslip and a coverslip coated with Sigmacote® (Sigma-Aldrich, SL2). The PAAm mixtures were prepared using different ratios of 40% acrylamide solution (Bio-Rad, 1610140) and 2% bis-acrylamide solution (Bio-Rad, 1610142) and Sigma molecular biology grade water to achieve different final stiffness, as previously described ^42^. Final water-PAAm mixtures were crosslinked with 0.5% APS and 0.15% N,N,N′,N′-tetramethylethylenediamine (ThermoFisher, 17919). 50 μL of the mixtures were added to each glass and allowed to crosslink for 30 minutes at room temperature. After crosslinking, Sigmacote®-treated coverslips were removed with a surgical blade and discarded. PAAm matrices were activated twice with 0.5 mg/mL Sulfo-SANPAH for 6 minutes at room temperature (Sigma-Aldrich, 80332) and coated with 5 μg/mL fibronectin (Sigma-Aldrich, F1141) for 1 hour at 37 ºC, then washed 3 times and preserved in PBS at 4 ºC no longer than 24 hours.

### Cell culture

RPE-1 cells were grown in Dulbecco’s Modified Eagle Medium: Nutrient Mixture F-12 (DMEM/F12, Lonza) supplemented with 10% Fetal Bovine Serum (GIBCO, Thermo Fisher Scientific), and 1% penicillin/streptomycin (Invitrogen). Cells were maintained at sub-confluency on standard cell culture plates and the medium was changed every 2-3 days. For cell mechanobiology experiments, RPE-1 cells were seeded at a confluency of 50% on hydrogel surfaces and were allowed to attach for approximately 12 hours in DMEM/F12 supplemented with 10% FBS and 1% penicillin/streptomycin before being processed. Cells were either preserved in RNAlater™ (ThermoFisher, AM7020) for RNA isolation or fixed in 4% paraformaldehyde (PFA) for immunostaining. For metabolic activity assays, cells were seeded at a confluency of 80% and cultured overnight.

### Cell metabolic activity

Cell medium was replaced by HBSS (ThermoScientific, 14175053) supplemented with 2% FBS. 5 mg/ml 3-[4,5-dimethylthiazol-2-yl]-2,5-diphenyl tetrazolium bromide (MTT) in RPMI-1640 was added to the cells in a volume equal to 10% of the culture volume. Cells were incubated at 37 ºC and 5% CO_2_ for 4 hours so MTT could be metabolized by mitochondrial dehydrogenases of viable cells, which cleave the tetrazolium ring yielding insoluble, purple MTT formazan crystals. Next, 0.1 N HCl in anhydrous isopropanol was added to the cells and used to solubilize any formazan crystals formed. Absorbance at 570 nm was measured and background absorbance at 690 nm was subtracted. Final absorbance signal is proportional to the number of metabolically active cells.

### Cytoskeleton inhibition

Drugs were added to cells 12 h after plating, followed by incubation for 1 h. Cytochalasin D (Sigma-Aldrich, C8273) and blebbistatin (Sigma-Aldrich, B0560) were added at a final concentration of 1 μM and 10 μM, respectively. After incubation cells were washed three times with PBS and prepared for immunostaining.

### Immunocytochemistry

Cells were washed three times with PBS and fixed with 4% PFA at room temperature (RT) for 10 min. Cells were then washed three times with PBS and permeabilized for 5 min with 0.2% Triton X-100 (Sigma-Aldrich) in 1% (w/v) BSA (Sigma-Aldrich). After permeabilization, samples were incubated with FBS for 1 hour at RT. Immunostaining with anti-YAP antibody took place at 4°C overnight. Secondary antibody, phalloidin and DAPI were incubated for 1 hour at RT. The following antibodies/reagents were used: YAP antibody (Santa Cruz Biotechnology YAP1 63.7; sc-101199), paxillin antibody (abcam; ab32084), DAPI (Invitrogen), Goat anti-Mouse IgG Alexa488 (Invitrogen), Goat anti-Mouse IgG Alexa568 (Invitrogen), Goat anti-Rabbit IgG Alexa594 (Invitrogen), Alexa647 phalloidin to stain actin (Invitrogen). Goat anti-Mouse IgG Alexa488 (Invitrogen) was used to stain YAP in all experiments except cytoskeleton inhibition ones, where Goat anti-Mouse IgG Alexa568 (Invitrogen) was employed to avoid green fluorescence of blebbistatin. Goat anti-Rabbit IgG Alexa594 (Invitrogen) was used to stain paxillin in focal adhesions. For measurement of YAP nuclear localization in the absence or presence of cytoskeleton inhibitors, images of DAPI/phalloidin/YAP antibody-stained cells were taken with a 40x NA 1.40 oil immersion objective using a laser scanning confocal microscope (Zeiss, LSM700), and analysed using ImageJ. YAP nuclear localization ratio was determined as the summed intensity of the YAP signal within a 10.8 μm^2^ area inside the nucleus divided by the summed intensity of the YAP signal within a 10.8 μm^2^ area inside the cytoplasm. To determine cell spreading, the outer membrane of cells was manually segmented and ImageJ was then used to compute total cell area. DAPI staining was used to quantify nuclei. Particles above 50 μm^2^ in size were analyzed. Length of focal adhesions was manually quantified on ImageJ.

### Retrograde flow measurements

To measure actin retrograde flow, cells were transfected with lifeact–GFP (Addgene plasmid 15238) using Lipofectamine 3000 transfection kit (Invitrogen, L3000008) 2 days before measurements. ∼12 hours before measurement, cells were plated on the protein hydrogels. Cells were imaged every second for 2 min with a a 20x W Plan-Apo DIC (UV) VIS-IR NA 1.0 water immersion dipping objective using a laser scanning confocal microscope (Zeiss, LSM780). For each cell, kymographs were obtained at the cell periphery, and actin speed was measured from the slope of actin features observed in the kymographs ^43^.

### Real-time quantitative PCR

Total RNA was extracted from cell samples using a combination of Trizol (ThermoFisher Scientific) and the RNAeasy micro kit (QIAGEN). For each sample, 150 ng RNA were reverse transcribed using the SuperScript™ IV VILO™ Master Mix (ThermoFisher Scientific). qPCR of cDNA was performed with SYBR Green (Applied Biosystems) accompanied by non-template controls. Results were normalized to endogenous GAPDH and β-Actin. Primer sequences can be found in **Supplementary Table 1**.

### Statistical analysis

In all figures, measurements are reported as mean ± standard error of the mean (SEM). The number of independent reproductions for each experiment is specified in the figure legends. Statistical significance was evaluated in GraphPad Prism. One-way ANOVA was used to assess differences between groups. When experimental data did not follow a normal distribution, Kruskal-Wallis test was applied instead. Differences were considered statistically significant at *p < 0.05, **p < 0.01, ***p < 0.001, and ****p < 0.0001.

## Supplementary Materials

- Supplementary Figures 1-11
- Supplementary Tables 1-2
- Supplementary Notes 1-3
- Supplementary References

