## Supplementary material for "Cell response to extracellular matrix energy dissipation outweighs rigidity sensing": HuertaLopez_SupplementaryInfo

**This PDF file includes:**

- 11 Supplementary Figures
- 2 Supplementary Tables
- 3 Supplementary Notes
- Supplementary References

(twitter: @AlegreCebollada)

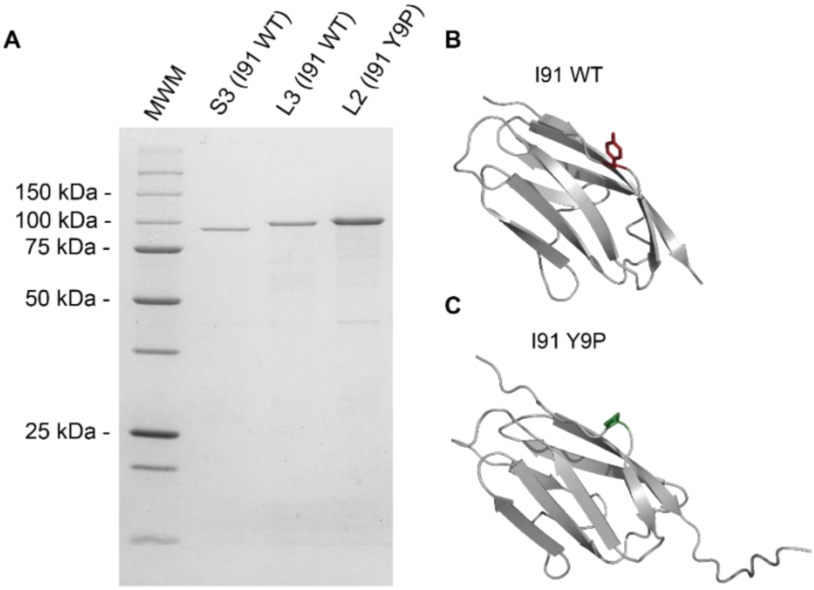


**Supplementary Figure 1: I91 polyproteins building blocks. (A)** 12 % SDS-PAGE analysis of representative purified soluble polyproteins used in this report. MWM: Precision Plus Protein Unstained Standards (Bio-Rad). **(B)** Three-dimensional structure of monomeric I91 (PDB: 1TIT [1]). The tyrosine residue employed to crosslink proteins is highlighted in red. **(C)** Three-dimensional structure of titin I91 Y9P domain (PDB: 2RQ8 [2]). Pro9 is shown in green.

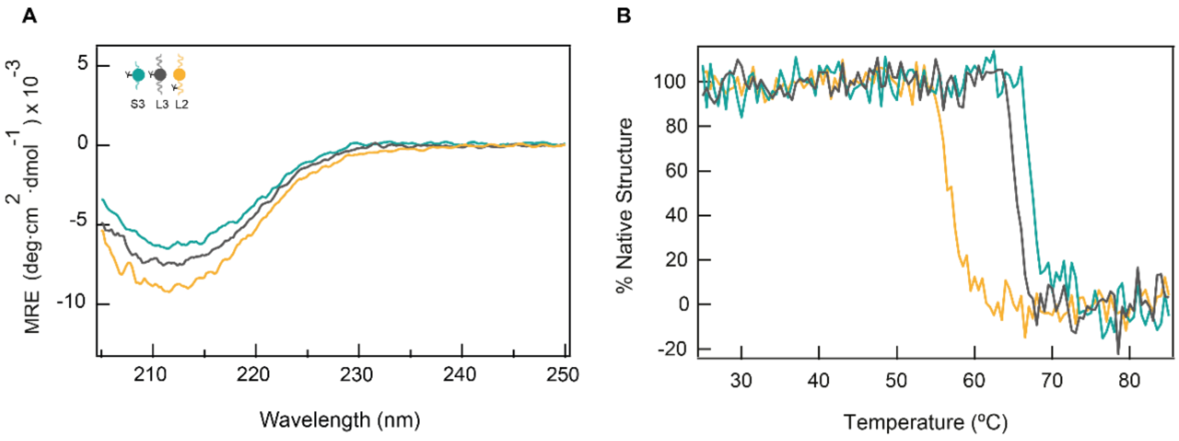


**Supplementary Figure 2: Thermal stability of protein building blocks by CD spectroscopy.** **(A)** CD spectra (given as mean residue ellipticity, MRE) obtained for S3 (teal), and L3 (grey) and L2 (mustard) domains recorded from 205 to 250 nm at 25 °C. Spectra show a minimum at ~212 nm, typical of the β–rich structure of the I91 [1]. **(B)** Thermal denaturation curves obtained by tracking the CD signal at 215 nm.

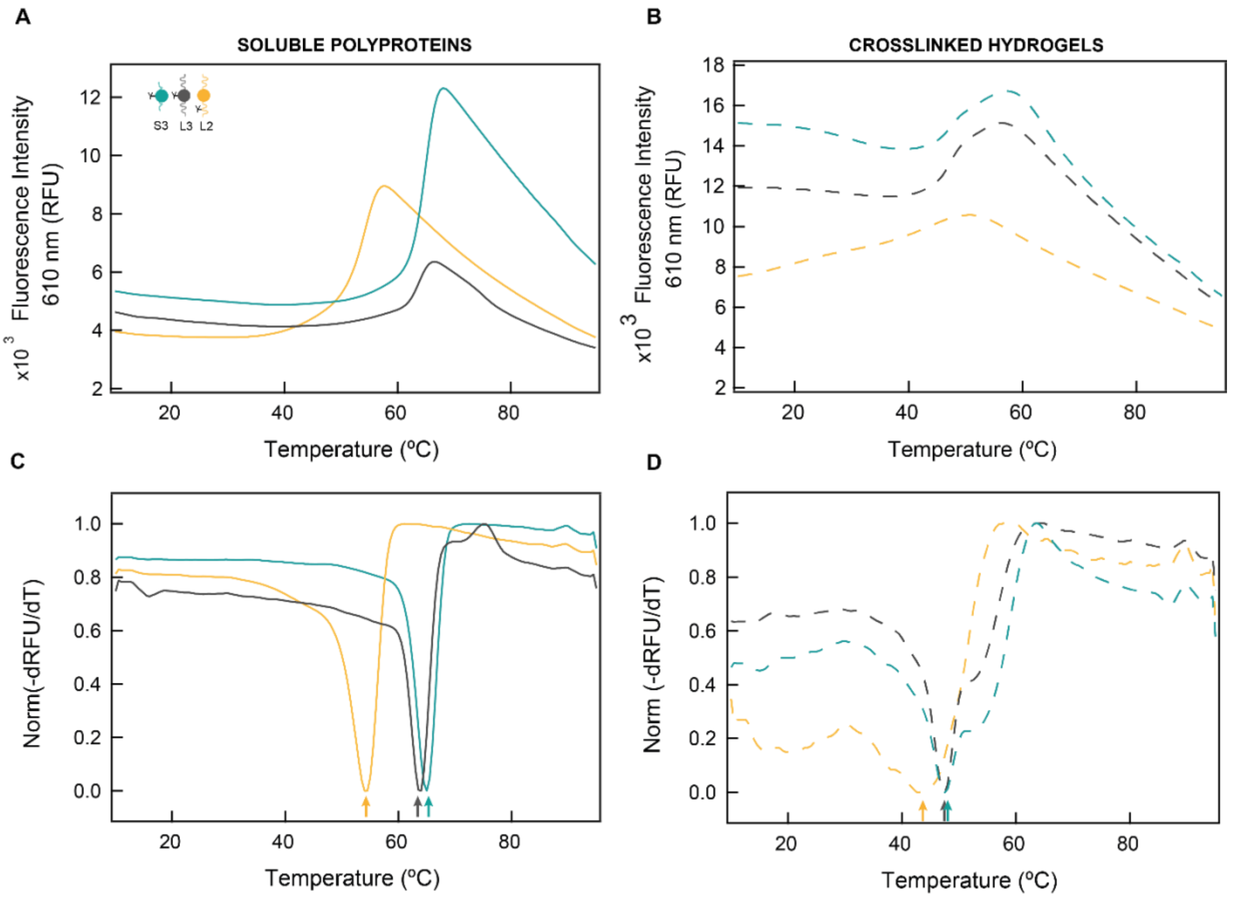


**Supplementary Figure 3: Polyprotein thermal stability by differential scanning fluorimetry (A, B)** Recording of fluorescence intensity versus temperature for the unfolding of soluble (A) I91 protein building blocks, and (B) I91protein hydrogels. **(C, D)** First derivative fluorescent signal of (C) I91 protein building blocks and (D) I91 protein hydrogels. Melting temperatures are indicated by arrows.

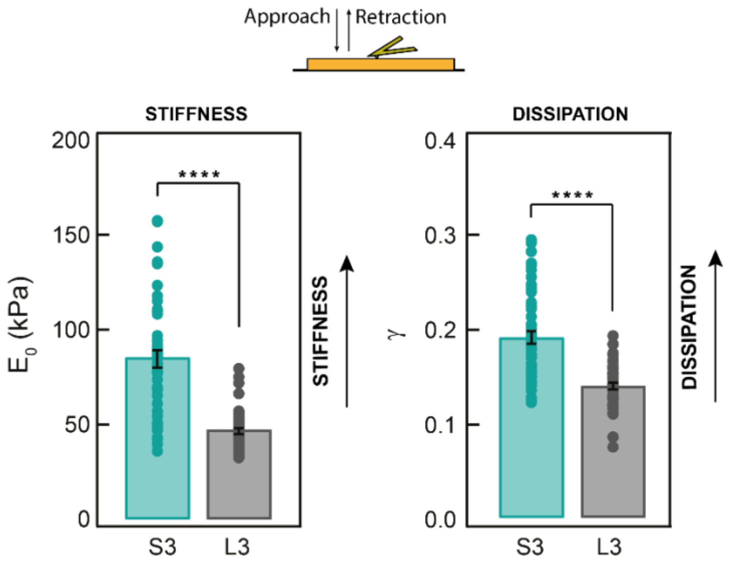


**Supplementary Figure 4: AFM analysis of the nanomechanics of I91-based hydrogels**. Hydrogels were assayed at 10 µm/s. Specimens were characterized after being incubated at 37 ºC for 12 hours. The two parameters of the power law rheology model, modulus $E_0$ (informing about stiffness) and exponent γ (related to energy dissipation), are shown. Data collected on 5 positions (10 determinations per position) from one hydrogel. Bars show mean ± SEM.

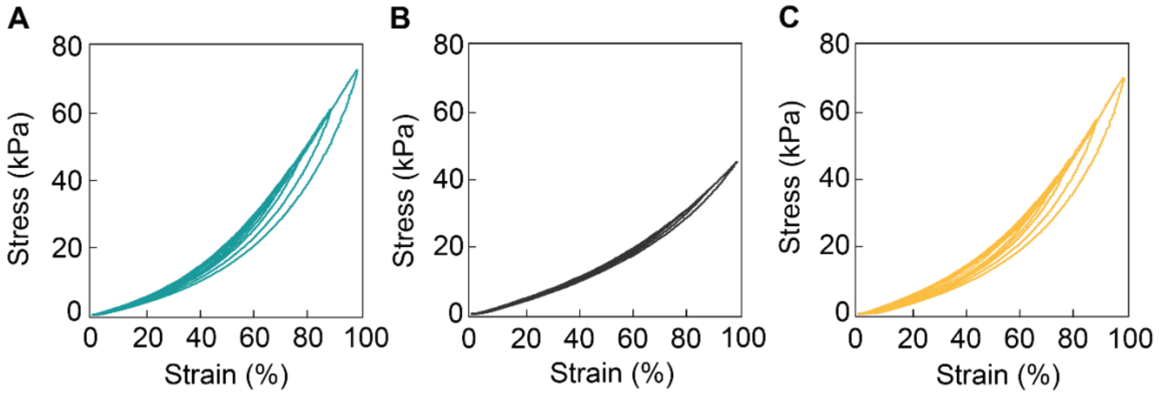


**Supplementary Figure 5: Tensile testing of I91 hydrogels**. Representative stress-strain curves of S3 **(A)**, L3 **(B)** and L2 **(C)** hydrogels loaded and unloaded at 5 mm/s. Curves were measured immediately one after another from low to high strains. These curves are also represented in **Fig. 1E,F** with an offset in the x-axis.

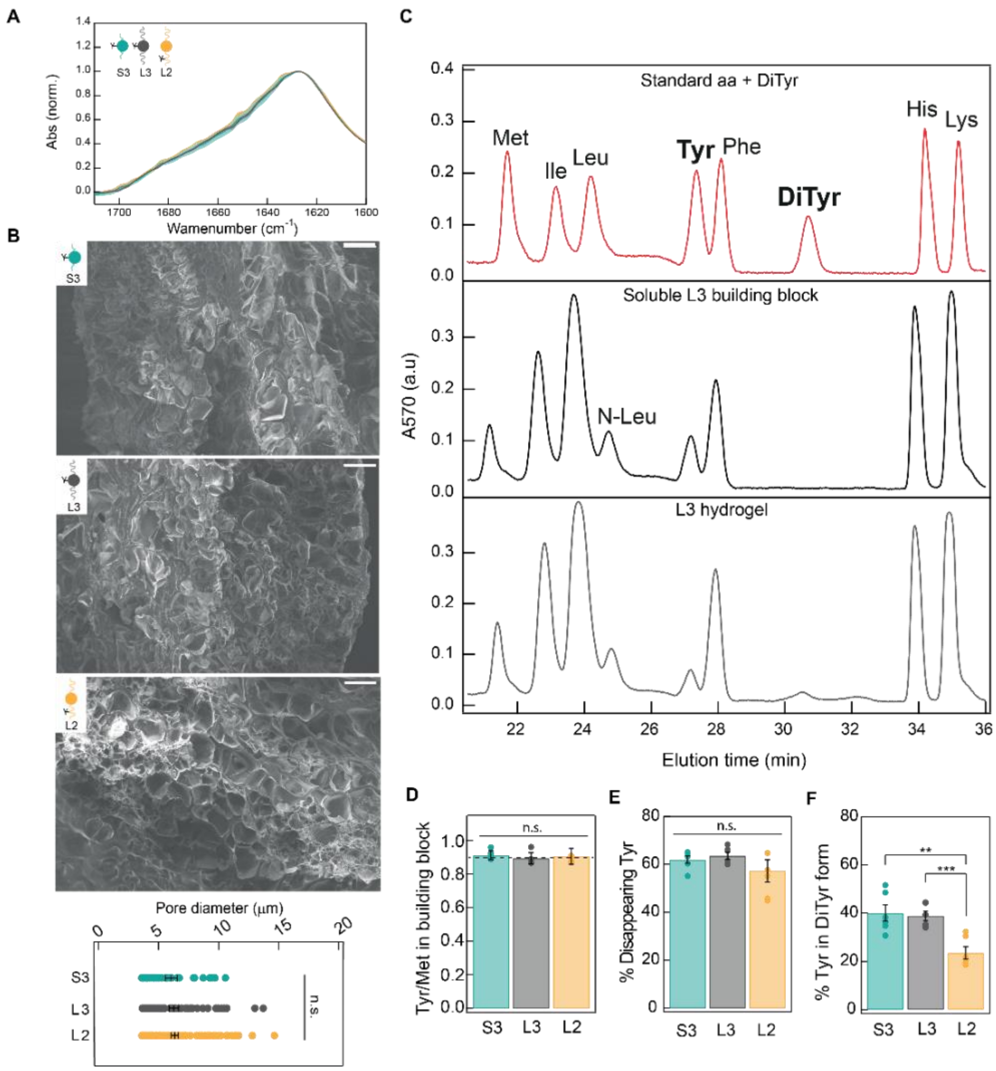


**Supplementary Fig. 6: Preservation of non-mechanical properties of S3, L3, and L2 hydrogels. (A)** Infrared spectra of S3, L3 and L2 hydrogels. Results represent the average value and SEM of three different determinations. Amide I bands of each spectrum are highly overlapping, indicating similar protein structures in the three hydrogels. Spectra were normalized according to the value at 1628 $cm^{-1}$. **(B)** Scanning electron microscopy images of S3 (top), L3 (center) and L2 (bottom) hydrogels. Images revealed the formation of a microporous network structure with pore sizes on the scale of a few micrometers, typical of protein hydrogels [3]. Scale bars are 10 µm. Bottom panel shows the quantification of S3, L3, and L2 pore size. Data were obtained from one EM image (45, 73, 103 pores for S3, L3 and L2 hydrogels, respectively). **(C)** Amino acids analysis of a mixture of protein amino acids and dityrosine standards (top, red). Amino acid analyses of soluble L3 polyprotein sample (middle, black) and of an L3 hydrogel (bottom, grey). Peaks corresponding to dityrosine are absent in the soluble polyprotein sample while they appear in samples coming from crosslinked hydrogels. **(D)** Experimental tyrosine to methionine ratios are equivalent for the three soluble building blocks and consistent with theoretical values (dotted line). **(E)** Percentage of tyrosines that disappear when soluble polyproteins are crosslinked into hydrogels. **(F)** Proportion of total tyrosines that appear as dityrosine in I91 hydrogels. Data in panels D-F represent mean ± SEM (n=6).

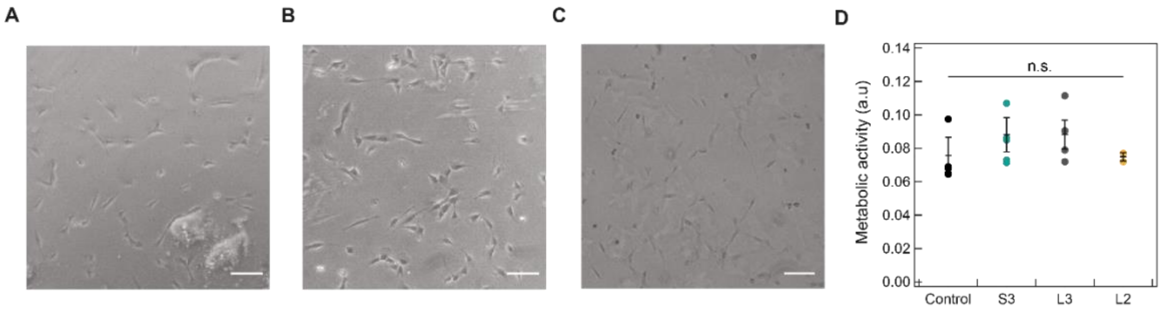


**Supplementary Figure 7: RPE-1 cells are equally metabolically active on S3, L3 and L2 hydrogels.** RPE-1 cells grown overnight on **(A)** S3, **(B)** L3 and **(C)** L2 protein hydrogels. **(D)** MTT assay of RPE-1 cells seeded on the three I91-based hydrogels. Cells grown on standard glass coverslips were used as control. Scale bars are 100 μm. Data are displayed as mean ± SEM (n=5 for all conditions except for L2 hydrogels, where n=2).

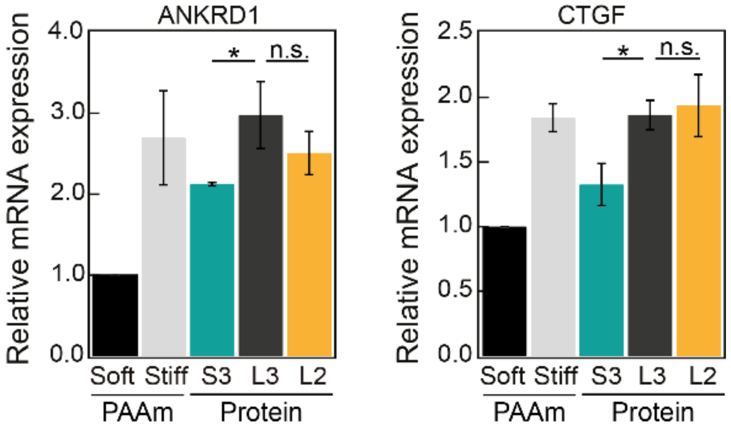


**Supplementary Figure 8: Expression of YAP target genes.** mRNA expression levels of YAP target genes ANKRD1 (left) and CTGF (right). n=3 independent experiments except for L2, where n=2.

57

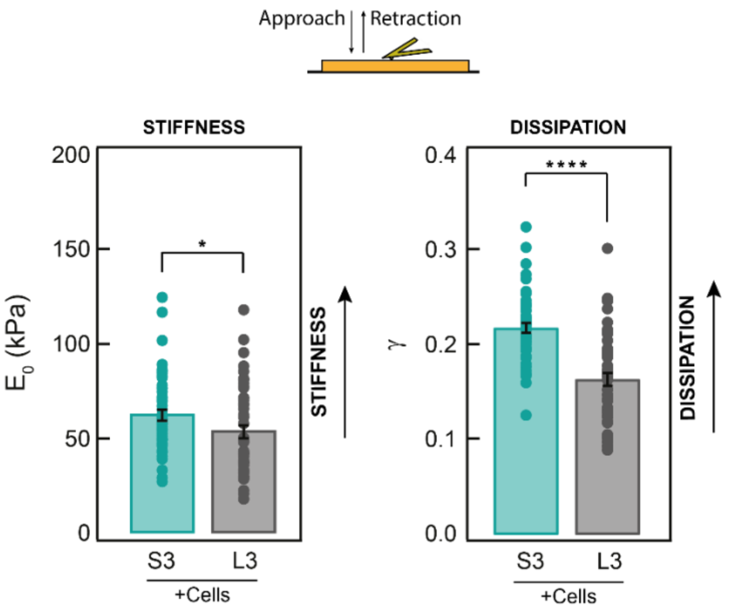


**Supplementary Figure 9: AFM analysis of the nanomechanics of cell-laden I91 hydrogels.** Hydrogels were assayed at 10 μm/s after 12 hours of being used as cell substrate at 37 ºC. The two parameters of the power law rheology model, modulus $E_0$ (informing about stiffness) and exponent γ (related to energy dissipation), are shown. Data collected on 5 positions (10 determinations per position) from one hydrogel. Bars show mean ± SEM.

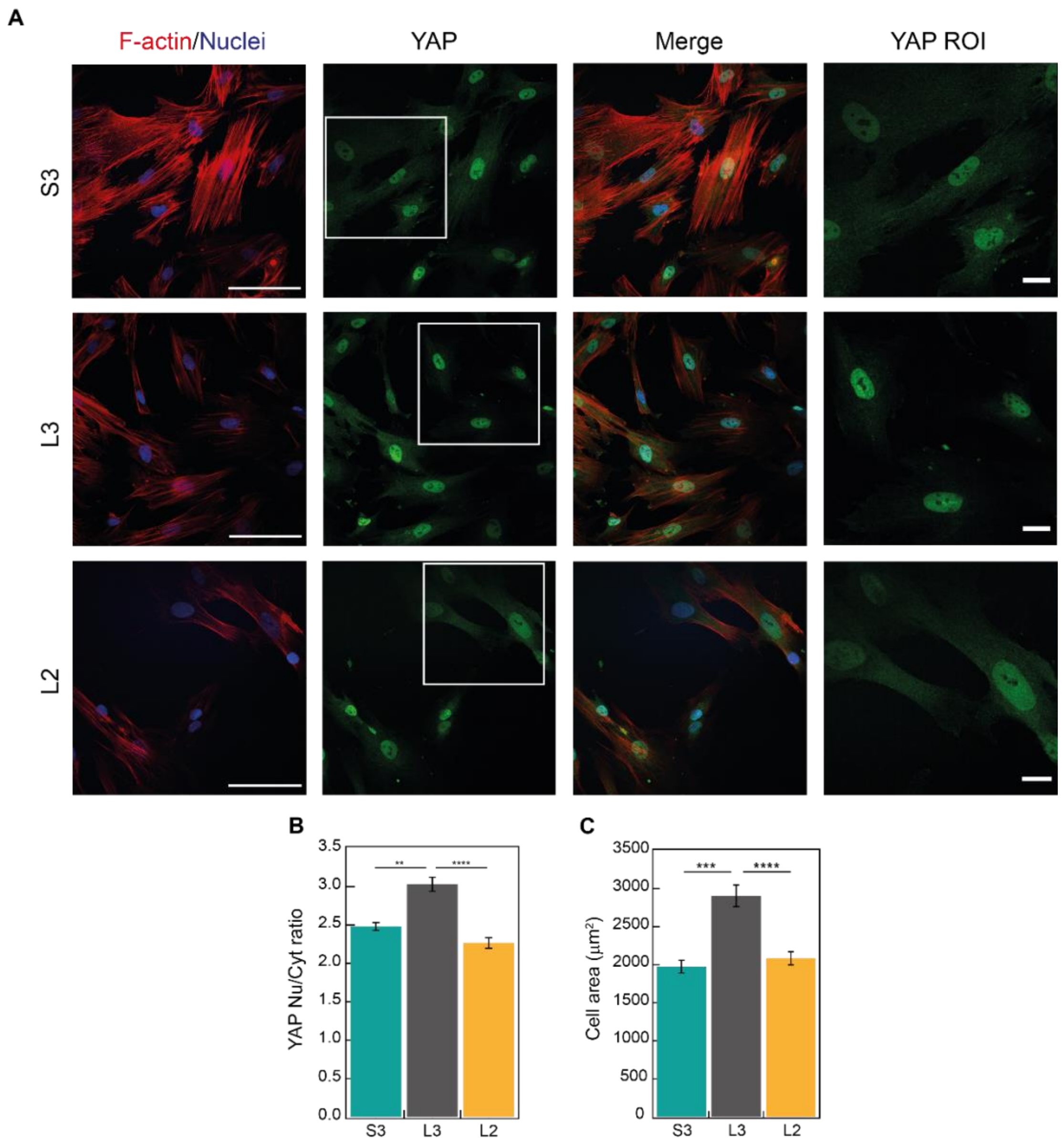


**Supplementary Figure 10: YAP activity in MSCs grown on viscoelastic protein hydrogel matrices. (A)** Confocal immunofluorescence images of YAP localization in MSCs grown on different I91 matrices. F-actin was stained with alexa647-conjugated phalloidin (red; left column), and nuclei were stained with DAPI (blue in merged images; first and third column). YAP was labelled with alexa488 conjugated antibody in green (second column). The column on the right shows zoomed views of the YAP ROI (boxed in white in the YAP images). Scale bars are 100 μm (left column) and 20 μm (right column). **(B)** Quantification of YAP cell distribution. **(C)** Cell spreading quantified as cell area. A minimum of n=30 cells per condition were quantified in a total of 3 independent experiments. Data are presented as mean ± SEM.

60

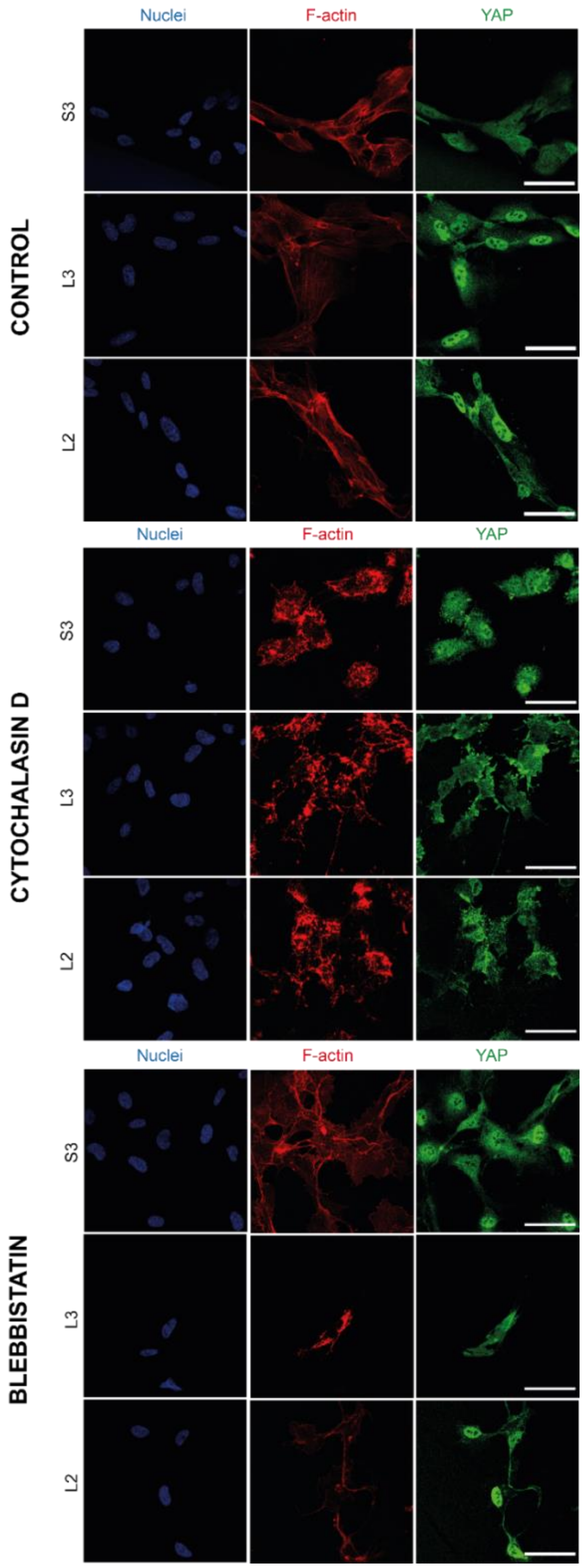


**Supplementary Figure 11: YAP localization in RPE-1 cells grown on viscoelastic protein hydrogel matrices is affected by treatment with actomyosin inhibitors.** Confocal immunofluorescence images of YAP localization in RPE-1 cells grown on different I91 matrices and treated with 1 μM cytochalasin D or 10 μM blebbistatin. Nuclei were stained with DAPI (blue; first column). F-actin was stained with alexa647-conjugated phalloidin (red; center column). YAP was labelled with alexa568-conjugated antibody (green; third column). Scale bars are 50 μm.

**Supplementary Table 1: Sequence of primers used in this study to add adapters to I91-Y9P domain or to follow mRNA expression levels of YAP target genes.**

| Primer | Sequence |
|---|---|
| **I_27 Y9P Fw** | 5'-CGCGGATCCGAGACCGTGCGTTTCCAGAGCCTAATAGAAGTGGAAAAGCCTCTGC -3' |
| **I_27 Y9P Rv** | 5'-ATAGGTACCTTAGCAACAAGATCTATAACCCAATTCTTTCACTTTCAGATTGGCT-3' |
| **CTGF Fw** | 5'-ACCGACTGGAAGACACGTTTG-3' |
| **CTGF Rv** | 5'-CCAGGTCAGCTTCGCAAGG-3' |
| **ANKRD 1 Fw** | 5'-AGTAGAGGAACTGGTCACTGG-3' |
| **ANKRD 1 Rv** | 5'-TGTTTCTCGCTTTTCCACTGTT-3' |
| **GAPDH Fw** | 5'-ATCACCATCTTCCAGGAGCG-3' |
| **GAPDH Rv** | 5'-CCTGCAAATGAGCCCCAG-3' |
| **β-Actin Fw** | 5'- CACCTTCCAGCAGATGTCGA-3' |
| **β-Actin Rv** | 5'-AGCATTTGCGGTGGACGATGG-3' |

**Supplementary Table 2: Parameters of I91 polyproteins**. Parameters were calculated according to ProtParam Tool [4].

| Name | Molecular weight (kDa) | Theoretical extinction coefficient ($g \cdot L^{-1} \cdot cm^{-1}$) |
|---|---|---|
| **S3** (I91 WT) | 81.476 | 0.686 |
| **L3** (I91 WT) | 89.413 | 0.625 |
| **L2** (I91 Y9P) | 89.493 | 0.625 |

**Supplementary Note 1. Sequence of $(I91)_8$ S3 recombinant construct.** Sequence including the start codon and a histidine tag used for protein purification is included. BamHI, BglII and BstYI restriction enzymes recognition sequences indicate ligation sites to the expression plasmid and modular assembly between domains, respectively.

BamHI-(I91-BstYI)$_8$-BglII

**<u>cDNA:</u>**

```
ATGAGAGGATCGCATCACCATCACCATCACGGATCCCTAATAGAAGTGGAAAAGCCTCTGTACGGAGT
AGAGGTGTTTGTTGGTGAAACAGCCCACTTTGAAATTGAACTTTCTGAACCTGATGTTCACGGCCAGT
GGAAGCTGAAAGGACAGCCTTTGGCAGCTTCCCCTGACTGTGAAATCATTGAGGATGGAAAGAAGCAT
ATTCTGATCCTTCATAACTGTCAGCTGGGTATGACAGGAGAGGTTTCCTTCCAGGCTGCTAATACCAA
ATCTGCAGCCAATCTGAAAGTGAAAGAATTGAGATCCCTAATAGAAGTGGAAAAGCCTCTGTACGGAG
TAGAGGTGTTTGTTGGTGAAACAGCCCACTTTGAAATTGAACTTTCTGAACCTGATGTTCACGGCCAG
TGGAAGCTGAAAGGACAGCCTTTGGCAGCTTCCCCTGACTGTGAAATCATTGAGGATGGAAAGAAGCA
TATTCTGATCCTTCATAACTGTCAGCTGGGTATGACAGGAGAGGTTTCCTTCCAGGCTGCTAATACCA
AATCTGCAGCCAATCTGAAAGTGAAAGAATTGAGATCCCTAATAGAAGTGGAAAAGCCTCTGTACGGA
GTAGAGGTGTTTGTTGGTGAAACAGCCCACTTTGAAATTGAACTTTCTGAACCTGATGTTCACGGCCA
GTGGAAGCTGAAAGGACAGCCTTTGGCAGCTTCCCCTGACTGTGAAATCATTGAGGATGGAAAGAAGC
ATATTCTGATCCTTCATAACTGTCAGCTGGGTATGACAGGAGAGGTTTCCTTCCAGGCTGCTAATACC
AAATCTGCAGCCAATCTGAAAGTGAAAGAATTGAGATCCCTAATAGAAGTGGAAAAGCCTCTGTACGG
AGTAGAGGTGTTTGTTGGTGAAACAGCCCACTTTGAAATTGAACTTTCTGAACCTGATGTTCACGGCC
AGTGGAAGCTGAAAGGACAGCCTTTGGCAGCTTCCCCTGACTGTGAAATCATTGAGGATGGAAAGAAG
CATATTCTGATCCTTCATAACTGTCAGCTGGGTATGACAGGAGAGGTTTCCTTCCAGGCTGCTAATAC
CAAATCTGCAGCCAATCTGAAAGTGAAAGAATTGAGATCCCTAATAGAAGTGGAAAAGCCTCTGTACG
GAGTAGAGGTGTTTGTTGGTGAAACAGCCCACTTTGAAATTGAACTTTCTGAACCTGATGTTCACGGC
CAGTGGAAGCTGAAAGGACAGCCTTTGGCAGCTTCCCCTGACTGTGAAATCATTGAGGATGGAAAGAA
GCATATTCTGATCCTTCATAACTGTCAGCTGGGTATGACAGGAGAGGTTTCCTTCCAGGCTGCTAATA
CCAAATCTGCAGCCAATCTGAAAGTGAAAGAATTGAGATCCCTAATAGAAGTGGAAAAGCCTCTGTAC
GGAGTAGAGGTGTTTGTTGGTGAAACAGCCCACTTTGAAATTGAACTTTCTGAACCTGATGTTCACGG
CCAGTGGAAGCTGAAAGGACAGCCTTTGGCAGCTTCCCCTGACTGTGAAATCATTGAGGATGGAAAGA
AGCATATTCTGATCCTTCATAACTGTCAGCTGGGTATGACAGGAGAGGTTTCCTTCCAGGCTGCTAAT
ACCAAATCTGCAGCCAATCTGAAAGTGAAAGAATTGAGATCCCTAATAGAAGTGGAAAAGCCTCTGTA
CGGAGTAGAGGTGTTTGTTGGTGAAACAGCCCACTTTGAAATTGAACTTTCTGAACCTGATGTTCACG
GCCAGTGGAAGCTGAAAGGACAGCCTTTGGCAGCTTCCCCTGACTGTGAAATCATTGAGGATGGAAAG
AAGCATATTCTGATCCTTCATAACTGTCAGCTGGGTATGACAGGAGAGGTTTCCTTCCAGGCTGCTAA
TACCAAATCTGCAGCCAATCTGAAAGTGAAAGAATTGAGATCCCTAATAGAAGTGGAAAAGCCTCTGT
ACGGAGTAGAGGTGTTTGTTGGTGAAACAGCCCACTTTGAAATTGAACTTTCTGAACCTGATGTTCAC
GGCCAGTGGAAGCTGAAAGGACAGCCTTTGGCAGCTTCCCCTGACTGTGAAATCATTGAGGATGGAAA
GAAGCATATTCTGATCCTTCATAACTGTCAGCTGGGTATGACAGGAGAGGTTTCCTTCCAGGCTGCTA
ATACCAAATCTGCAGCCAATCTGAAAGTGAAAGAATTGAGATCTTAA
```

**Protein:**

MRGSHHHHHHGSLIEVEKPLYGVEVFVGETAHFEIELSEPDVHGQWKLKGQPLAASPDCEIIEDGKKHILILHNCQLGMTGEVSFQAANTKSAANLKVKELRSLIEVEKPLYGVEVFVGETAHFEIELSEPDVHGQWKLKGQPLAASPDCEIIEDGKKHILILHNCQLGMTGEVSFQAANTKSAANLKVKELRSLIEVEKPLYGVEVFVGETAHFEIELSEPDVHGQWKLKGQPLAASPDCEIIEDGKKHILILHNCQLGMTGEVSFQAANTKSAANLKVKELRSLIEVEKPLYGVEVFVGETAHFEIELSEPDVHGQWKLKGQPLAASPDCEIIEDGKKHILILHNCQLGMTGEVSFQAANTKSAANLKVKELRSLIEVEKPLYGVEVFVGETAHFEIELSEPDVHGQWKLKGQPLAASPDCEIIEDGKKHILILHNCQLGMTGEVSFQAANTKSAANLKVKELRSLIEVEKPLYGVEVFVGETAHFEIELSEPDVHGQWKLKGQPLAASPDCEIIEDGKKHILILHNCQLGMTGEVSFQAANTKSAANLKVKELRSLIEVEKPLYGVEVFVGETAHFEIELSEPDVHGQWKLKGQPLAASPDCEIIEDGKKHILILHNCQLGMTGEVSFQAANTKSAANLKVKELRS

**Supplementary Note 2. Sequence of (I91)$_8$ L3 recombinant construct.** Sequence including the start codon providing and a histidine tag used for protein purification is included. BamHI, BglII and BstYI restriction enzymes recognition sequences indicate ligation sites to the expression plasmid and modular assembly between domains, respectively.

BamHI-(linker-I91-GlySer-BstYI)$_8$-BglII

**cDNA:**

```
ATGAGAGGATCGCATCACCATCACCATCACGGATCCGAGACCGTGCGTTTCCAGAGCCTAATAGAAGT
GGAAAAGCCTCTGTACGGAGTAGAGGTGTTTGTTGGTGAAACAGCCCACTTTGAAATTGAACTTTCTG
AACCTGATGTTCACGGCCAGTGGAAGCTGAAAGGACAGCCTTTGGCAGCTTCCCCTGACTGTGAAATC
ATTGAGGATGGAAAGAAGCATATTCTGATCCTTCATAACTGTCAGCTGGGTATGACAGGAGAGGTTTC
CTTCCAGGCTGCTAATACCAAATCTGCAGCCAATCTGAAAGTGAAAGAATTGGGTTCTAGATCCGAAA
CCGTTCGCTTTCAGTCACTGATTGAAGTTGAAAAACCGCTGTATGGTGTGGAAGTTTTTGTGGGCGAA
ACCGCACATTTTGAAATCGAGCTGAGCGAACCGGATGTGCATGGTCAATGGAAACTGAAGGGTCAGCC
GCTGGCAGCAAGTCCGGATTGCGAAATTATTGAAGATGGCAAAAAACACATCCTGATTCTGCATAATT
GCCAACTGGGCATGACCGGTGAAGTTAGCTTTCAGGCAGCAAATACAAAAAGCGCTGCAAACCTGAAG
GTTAAAGAACTGGGTAGCCGTAGCGAAACAGTACGTTTTCAGAGTCTGATCGAGGTGGAAAAACCGTT
ATATGGCGTTGAGGTTTTCGTAGGTGAGACTGCCCATTTCGAGATCGAACTGTCAGAGCCAGATGTAC
ATGGACAGTGGAAGTTAAAAGGCCAACCACTGGCAGCATCACCAGATTGTGAGATCATCGAGGACGGT
AAAAAACACATTCTTATCCTGCACAACTGTCAATTAGGTATGACTGGCGAAGTTTCATTTCAAGCAGC
CAATACGAAATCAGCCGCTAACTTAAAAGTAAAAGAGCTGGGTTCACGTTCAGAAACGGTTCGTTTTC
AAAGCCTTATCGAAGTAGAGAAACCACTGTACGGCGTTGAAGTATTCGTTGGAGAAACGGCTCATTTT
GAGATTGAGTTAAGTGAGCCGGACGTTCACGGACAATGGAAATTAAAGGGACAACCGTTAGCCGCTTC
TCCCGATTGTGAAATAATCGAAGATGGGAAGAAACACATTTTGATCTTACACAATTGCCAGTTAGGGA
TGACAGGGGAAGTGAGTTTTCAAGCCGCAAACACCAAAAGTGCAGCGAATTTAAAGGTGAAAGAGTTA
GGTAGCCGCTCAGAAACAGTGAGATTTCAGAGCTTAATTGAGGTTGAGAAACCCCTTTATGGCGTCGA
GGTCTTTGTCGGCGAGACAGCACACTTCGAGATTGAATTATCAGAACCCGACGTGCATGGCCAGTGGA
AACTTAAAGGGCAACCTCTTGCAGCCAGTCCAGACTGCGAGATAATAGAGGACGGCAAGAAGCACATA
TTAATCTTGCATAATTGTCAGCTTGGAATGACTGGTGAAGTGTCGTTCCAGGCAGCGAACACTAAATC
AGCTGCAAATTTGAAAGTCAAAGAACTTGGCAGCCGTTCTGAAACTGTGCGCTTCCAATCTCTTATTG
AGGTAGAAAAGCCGCTTTACGGTGTCGAAGTGTTCGTGGGTGAGACAGCGCATTTTGAAATAGAATTG
TCAGAACCGGATGTACACGGCCAATGGAAGTTAAAGGGTCAGCCGCTTGCCGCATCACCGGACTGTGA
GATTATAGAAGATGGTAAAAAGCATATCTTAATTCTTCACAACTGCCAGCTTGGCATGACTGGCGAGG
TGAGTTTTCAGGCTGCGAATACTAAGAGCGCAGCGAATCTGAAGGTAAAAGAGCTTGGCTCTCGTAGC
GAAACGGTTCGCTTCCAGAGTTTAATTGAAGTCGAGAAGCCGTTATACGGGGTAGAAGTCTTTGTGGG
AGAAACTGCGCACTTTGAGATAGAACTGAGTGAACCAGACGTACACGGTCAGTGGAAGCTTAAGGGGC
AGCCGTTAGCAGCGAGCCCTGATTGCGAGATTATCGAGGATGGGAAAAAGCACATACTGATTTTACAC
AACTGTCAACTGGGAATGACAGGGGAAGTGTCATTTCAAGCGGCAAATACTAAAAGTGCCGCAAATCT
TAAAGTAAAAGAATTAGGTAGTCGCAGCGAAACCGTCAGATTCCAAAGCCTGATAGAGGTCGAGAAGC
CCCTGTATGGGGTTGAAGTGTTCGTAGGCGAAACAGCTCACTTCGAAATCGAGTTATCCGAGCCGGAT
GTCCACGGTCAGTGGAAATTGAAAGGTCAGCCATTAGCAGCGTCACCCGATTGCGAAATCATAGAGGA
TGGGAAAAAACACATCTTAATATTGCATAACTGCCAATTAGGAATGACAGGTGAGGTTAGCTTCCAAG
CGGCAAACACGAAATCCGCTGCCAACTTGAAGGTGAAAGAATTAGGCAGCAGATCTTAA
```

**Protein:**

MRGSHHHHHHGSETVRFQSLIEVEKPLYGVEVFVGETAHFEIELSEPDVHGQWKLKGQPLAASPDCEI
IEDGKKHILILHNCQLGMTGEVSFQAANTKSAANLKVKELGSRSETVRFQSLIEVEKPLYGVEVFVGE
TAHFEIELSEPDVHGQWKLKGQPLAASPDCEIIEDGKKHILILHNCQLGMTGEVSFQAANTKSAANLK
VKELGSRSETVRFQSLIEVEKPLYGVEVFVGETAHFEIELSEPDVHGQWKLKGQPLAASPDCEIIEDG
KKHILILHNCQLGMTGEVSFQAANTKSAANLKVKELGSRSETVRFQSLIEVEKPLYGVEVFVGETAHF
EIELSEPDVHGQWKLKGQPLAASPDCEIIEDGKKHILILHNCQLGMTGEVSFQAANTKSAANLKVKEL
GSRSETVRFQSLIEVEKPLYGVEVFVGETAHFEIELSEPDVHGQWKLKGQPLAASPDCEIIEDGKKHI
LILHNCQLGMTGEVSFQAANTKSAANLKVKELGSRSETVRFQSLIEVEKPLYGVEVFVGETAHFEIEL
SEPDVHGQWKLKGQPLAASPDCEIIEDGKKHILILHNCQLGMTGEVSFQAANTKSAANLKVKELGSRS
ETVRFQSLIEVEKPLYGVEVFVGETAHFEIELSEPDVHGQWKLKGQPLAASPDCEIIEDGKKHILILH
NCQLGMTGEVSFQAANTKSAANLKVKELGSRSETVRFQSLIEVEKPLYGVEVFVGETAHFEIELSEPD
VHGQWKLKGQPLAASPDCEIIEDGKKHILILHNCQLGMTGEVSFQAANTKSAANLKVKELGSRS

**Supplementary Note 3. Sequence of (I91-Y9P)$_8$ L2 recombinant construct.** Sequence including the start codon and a histidine tag used for protein purification is included. BamHI, BglII and BstYI restriction enzymes recognition sequences indicate ligation sites to the expression plasmid and modular assembly between domains, respectively.

BamHI-(linker-I91Y9P-GlyTyr-BstYI)$_8$-BglII

**cDNA:**

ATGAGAGGATCGCATCACCATCACCATCACGGATCCGAGACCGTGCGTTTCCAGAGCCTAATAGAAGT
GGAAAAGCCTCTGCCGGGAGTAGAGGTGTTTGTTGGTGAAACAGCCCACTTTGAAATTGAACTTTCTG
AACCTGATGTTCACGGCCAGTGGAAGCTGAAAGGACAGCCTTTGGCAGCTTCCCCTGACTGTGAAATC
ATTGAGGATGGAAAGAAGCATATTCTGATCCTTCATAACTGTCAGCTGGGTATGACAGGAGAGGTTTC
CTTCCAGGCTGCTAATACCAAATCTGCAGCCAATCTGAAAGTGAAAGAATTGGGTTATAGATCCGAGA
CCGTGCGTTTCCAGAGCCTAATAGAAGTGGAAAAGCCTCTGCCGGGAGTAGAGGTGTTTGTTGGTGAA
ACAGCCCACTTTGAAATTGAACTTTCTGAACCTGATGTTCACGGCCAGTGGAAGCTGAAAGGACAGCC
TTTGGCAGCTTCCCCTGACTGTGAAATCATTGAGGATGGAAAGAAGCATATTCTGATCCTTCATAACT
GTCAGCTGGGTATGACAGGAGAGGTTTCCTTCCAGGCTGCTAATACCAAATCTGCAGCCAATCTGAAA
GTGAAAGAATTGGGTTATAGATCCGAGACCGTGCGTTTCCAGAGCCTAATAGAAGTGGAAAAGCCTCT
GCCGGGAGTAGAGGTGTTTGTTGGTGAAACAGCCCACTTTGAAATTGAACTTTCTGAACCTGATGTTC
ACGGCCAGTGGAAGCTGAAAGGACAGCCTTTGGCAGCTTCCCCTGACTGTGAAATCATTGAGGATGGA
AAGAAGCATATTCTGATCCTTCATAACTGTCAGCTGGGTATGACAGGAGAGGTTTCCTTCCAGGCTGC
TAATACCAAATCTGCAGCCAATCTGAAAGTGAAAGAATTGGGTTATAGATCCGAGACCGTGCGTTTCC
AGAGCCTAATAGAAGTGGAAAAGCCTCTGCCGGGAGTAGAGGTGTTTGTTGGTGAAACAGCCCACTTT
GAAATTGAACTTTCTGAACCTGATGTTCACGGCCAGTGGAAGCTGAAAGGACAGCCTTTGGCAGCTTC
CCCTGACTGTGAAATCATTGAGGATGGAAAGAAGCATATTCTGATCCTTCATAACTGTCAGCTGGGTA
TGACAGGAGAGGTTTCCTTCCAGGCTGCTAATACCAAATCTGCAGCCAATCTGAAAGTGAAAGAATTG
GGTTATAGATCCGAGACCGTGCGTTTCCAGAGCCTAATAGAAGTGGAAAAGCCTCTGCCGGGAGTAGA
GGTGTTTGTTGGTGAAACAGCCCACTTTGAAATTGAACTTTCTGAACCTGATGTTCACGGCCAGTGGA
AGCTGAAAGGACAGCCTTTGGCAGCTTCCCCTGACTGTGAAATCATTGAGGATGGAAAGAAGCATATT
CTGATCCTTCATAACTGTCAGCTGGGTATGACAGGAGAGGTTTCCTTCCAGGCTGCTAATACCAAATC
TGCAGCCAATCTGAAAGTGAAAGAATTGGGTTATAGATCCGAGACCGTGCGTTTCCAGAGCCTAATAG
AAGTGGAAAAGCCTCTGCCGGGAGTAGAGGTGTTTGTTGGTGAAACAGCCCACTTTGAAATTGAACTT
TCTGAACCTGATGTTCACGGCCAGTGGAAGCTGAAAGGACAGCCTTTGGCAGCTTCCCCTGACTGTGA
AATCATTGAGGATGGAAAGAAGCATATTCTGATCCTTCATAACTGTCAGCTGGGTATGACAGGAGAGG
TTTCCTTCCAGGCTGCTAATACCAAATCTGCAGCCAATCTGAAAGTGAAAGAATTGGGTTATAGATCC
GAGACCGTGCGTTTCCAGAGCCTAATAGAAGTGGAAAAGCCTCTGCCGGGAGTAGAGGTGTTTGTTGG
TGAAACAGCCCACTTTGAAATTGAACTTTCTGAACCTGATGTTCACGGCCAGTGGAAGCTGAAAGGAC
AGCCTTTGGCAGCTTCCCCTGACTGTGAAATCATTGAGGATGGAAAGAAGCATATTCTGATCCTTCAT
AACTGTCAGCTGGGTATGACAGGAGAGGTTTCCTTCCAGGCTGCTAATACCAAATCTGCAGCCAATCT
GAAAGTGAAAGAATTGGGTTATAGATCCGAGACCGTGCGTTTCCAGAGCCTAATAGAAGTGGAAAAGC
CTCTGCCGGGAGTAGAGGTGTTTGTTGGTGAAACAGCCCACTTTGAAATTGAACTTTCTGAACCTGAT
GTTCACGGCCAGTGGAAGCTGAAAGGACAGCCTTTGGCAGCTTCCCCTGACTGTGAAATCATTGAGGA
TGGAAAGAAGCATATTCTGATCCTTCATAACTGTCAGCTGGGTATGACAGGAGAGGTTTCCTTCCAGG
CTGCTAATACCAAATCTGCAGCCAATCTGAAAGTGAAAGAATTGGGTTATAGATCTTAA

**Protein:**

MRGSHHHHHHGSETVRFQSLIEVEKPLPGVEVFVGETAHFEIELSEPDVHGQWKLKGQPLAASPDCEI
IEDGKKHILILHNCQLGMTGEVSFQAANTKSAANLKVKELGYRSETVRFQSLIEVEKPLPGVEVFVGE
TAHFEIELSEPDVHGQWKLKGQPLAASPDCEIIEDGKKHILILHNCQLGMTGEVSFQAANTKSAANLK
VKELGYRSETVRFQSLIEVEKPLPGVEVFVGETAHFEIELSEPDVHGQWKLKGQPLAASPDCEIIEDG
KKHILILHNCQLGMTGEVSFQAANTKSAANLKVKELGYRSETVRFQSLIEVEKPLPGVEVFVGETAHF
EIELSEPDVHGQWKLKGQPLAASPDCEIIEDGKKHILILHNCQLGMTGEVSFQAANTKSAANLKVKEL
GYRSETVRFQSLIEVEKPLPGVEVFVGETAHFEIELSEPDVHGQWKLKGQPLAASPDCEIIEDGKKHI
LILHNCQLGMTGEVSFQAANTKSAANLKVKELGYRSETVRFQSLIEVEKPLPGVEVFVGETAHFEIEL
SEPDVHGQWKLKGQPLAASPDCEIIEDGKKHILILHNCQLGMTGEVSFQAANTKSAANLKVKELGYRS
ETVRFQSLIEVEKPLPGVEVFVGETAHFEIELSEPDVHGQWKLKGQPLAASPDCEIIEDGKKHILILH
NCQLGMTGEVSFQAANTKSAANLKVKELGYRSETVRFQSLIEVEKPLPGVEVFVGETAHFEIELSEPD
VHGQWKLKGQPLAASPDCEIIEDGKKHILILHNCQLGMTGEVSFQAANTKSAANLKVKELGYRS
